## Supplementary Material for "Sustained Enzymatic Activity and Flow in Crowded Protein Droplets"

(Dated: May 15, 2021)

### SUPPLEMENTAL MATERIALS

#### Materials

Polyethylene glycol (PEG) 4000 Da (A16151) was purchased from Alpha-Aesar. Bovine serum albumin (BSA) (A7638), Potassium phospate dibasic trihydrate (60349), 3-(Trimethoxysilyl)propylmethacrylate (440159), 2-Hydroxy-4'-(2-hydroxyyethoxy)-2-methylpropiophenone (410896), Poly(ethylene glycol) diacrylate (PEGDA) 700 Da (455008), L-lactate dehydrogenase (LDH) from muscle rabbit (L1254), Jack bean urease (U4002), $\beta$-Nicotinamide adenine dinucleotide, reduced disodium salt hydrate (NADH) (N8129), Pyruvate (P8524), Lactate assay Kit (MAK064), Phenol nitroprusside solution (P6994), Alkaline hypochlorite solution (A1727), Urease activity Assay Kit (MAK120), Rhodamine-B (R6626) were purchased from Sigma-Aldrich. Potassium phosphate monobasic (42420) was purchased from Acros Organics. Deuterium oxide ($D_2O$) (DE50B) was purchased from Apollo. N,N-dimethylformamide (DMF) (D119) was purchased from Fisher Chemicals. Alexa Fluor 594 and 488 Microscale Protein Labeling Kit (A30008 and A30006) and Carboxylic acid, Acetate, Succinimidyl Ester SNARF-1 (S2280) were purchased from Thermo Fischer Scientific. Trichloroacetic acid (TCA) (34603) was purchased from Nacali Tesque. Fluorescent polystyrene nanoparticles tracers of 0.2 and 1 $\mu$m of diameter (FCDG003, FCDG006) were purchased from Bangs Laboratories. Fluorescein isothiocyanate (FITC) (AB178737) was purchased from abcr.

#### Droplet formation

A polyethylene glycol stock solution at 600 mg/mL was prepared mixing the appropriate amount of PEG 4000 Da and Milli-Q water. The pH of the solution was measured to be 7 with a pH meter (Orion Star A111, Thermo Fisher and F71, HORIBA Scientific).

A bovine serum albumin stock solution with a target concentration of 5 mM (332 mg/mL) was prepared mixing the appropriate amount of BSA and Milli-Q water. The actual concentration was confirmed measuring the absorption intensity at the 280 nm peak with a UV-vis spectrophotometer (Cary 60 Spectrophotometer, Agilent Technologies and UV-1900 UV-Vis Spectrophotometer, Shimadzu), assuming $\varepsilon_{280} = 43824\ \mathrm{M^{-1}cm^{-1}}$ (https://web.expasy.org/protparam/). The pH of the solution was measured to be 7.

A potassium phosphate buffer (KP) stock solution at 500 mM and pH 7 was prepared.

The droplet suspensions were prepared mixing appropriate amounts of PEG, BSA stocks, together with KP buffer, KCl and Milli-Q water in an Eppendorf tube. At the standard working conditions the final concentrations of the components were 230 mg/mL of PEG, 30 mg/mL BSA, 200 mM KCl and 100 mM KP pH 7.0. Additional components like enzymes, cofactors, substrates and fluorescent nanoparticles were also added to the tube when needed. The tubes were gently shaken by hand to promote the mixing. When required, the supernatant was isolated centrifuging the droplet suspension for 30 minutes at 16900 g at room temperature and extracting the top phase with a pipette.

#### Phase diagram

The phase diagram was mapped as a function of BSA and PEG composition. Specifically, we determined the two arms of the binodal by measuring the concentrations of BSA and PEG in both droplet and supernatant phases for various average sample compositions. All samples were prepared and measured in triplicate at room temperature. The two phases were obtained by centrifuging the phase-separated samples for 60 min at 16900 g and then carefully transferring the supernatant to a separate tube using a micropipette. The centrifugation step was repeated and any residual supernatant removed from the droplet phase.

The BSA concentrations in the two phases were determined by measuring absorbance at 280 nm as explained in Supplementary Materials section. Measurements were performed in a quartz cuvette (UQ-124, Portmann Technologies) on samples diluted with Milli-Q water (typically 200-fold diluted for droplet phase and 10- to 40-fold diluted for supernatant phase).

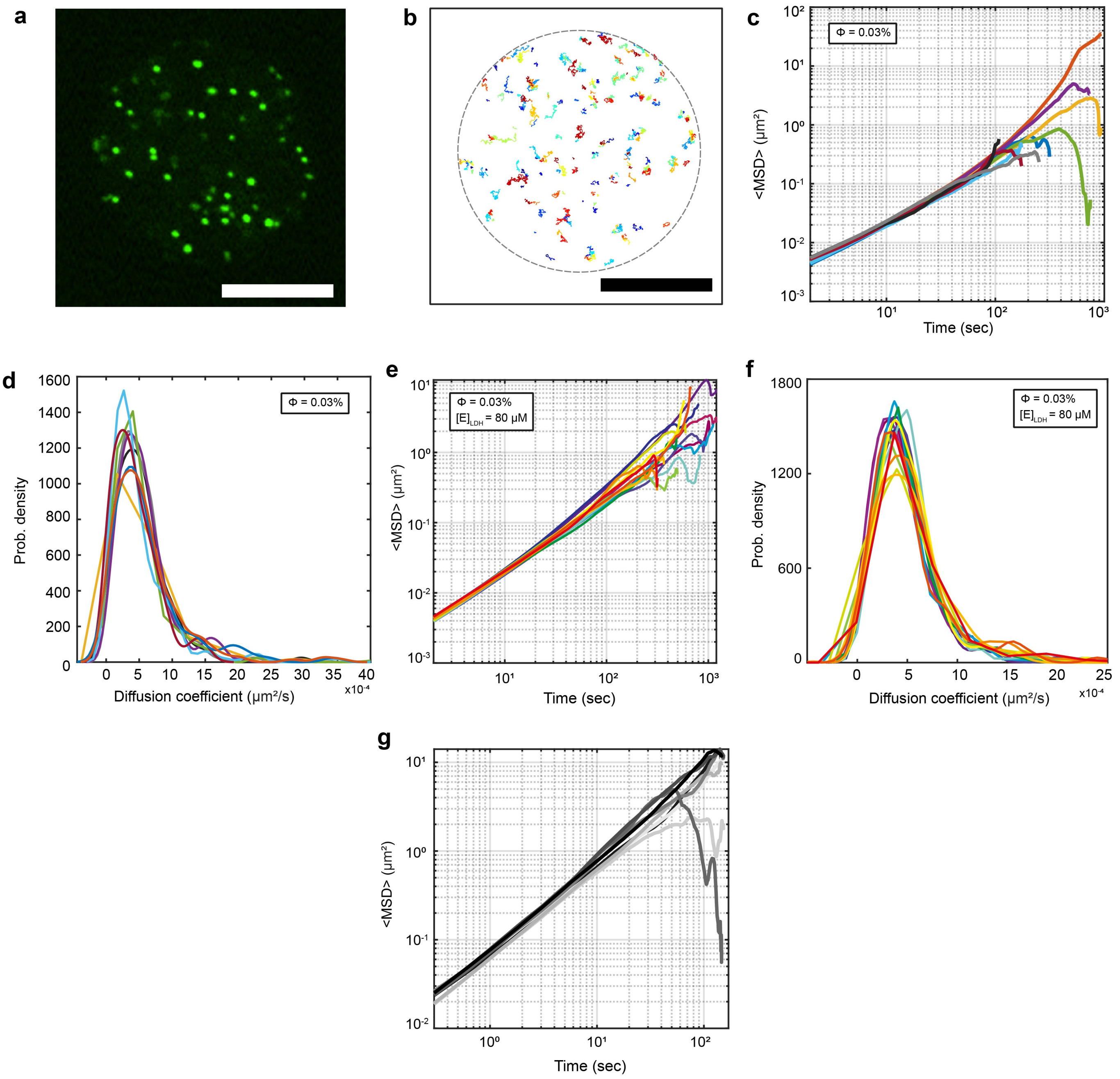


FIG. S1. **Supplementary Figure: Single particle tracking experiments.** a, Confocal microscopy image of fluorescent tracers of 200 nm radius partitioned in a BSA droplet. Scalebar = 10 $\mu$m. b, Trajectories of the fluorescent particles for the droplet in panel a, over a period of 20 minutes. The gray dashed line is the droplet outline. Scalebar = 10 $\mu$m. c, Mean squared displacement versus time for seven different droplets at the chosen working conditions and d, corresponding diffusion coefficient kernel density distribution. e, Mean squared displacement versus time for several different droplets containing LDH and f, corresponding diffusion coefficient kernel density distribution. g, Mean squared displacement versus time for ten different experiments for particles dispersed in supernatant.

PEG concentrations were determined using proton nuclear magnetic resonance ($^1$H NMR) spectroscopy [86] with $D_2O$ as solvent. DMF was added to the sample at an appropriate final concentration (10 - 660 mM) to serve as a calibration standard for the PEG concentration measurement. Measurements were performed in disposable 5 mm NMR tubes (Type E, Schott) on samples diluted with $D_2O$ (20-fold diluted for droplet phase and 200-fold diluted for supernatant phase). $^1$H NMR spectra were acquired using a 300 MHz instrument (300 Ultrashield, Bruker) at 298 K and averaged 16 times The resulting data were analyzed using the MestreNova soft-

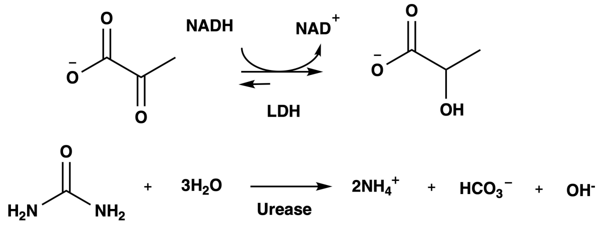


FIG. S2. **Supplementary Figure: Scheme of the chemical reactions performed by LDH and urease.**

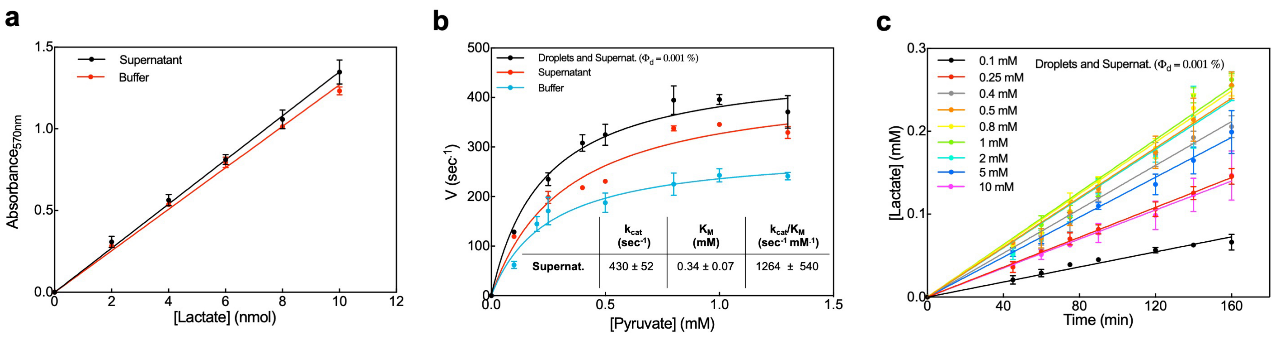


FIG. S3. **Supplementary Figure: Michaelis-Menten kinetics of LDH in droplets and supernatant and LDH free in the supernatant phase and in buffer.** a, Calibration curve of lactate in buffer (red) and in the supernatant (black). Lactate from the Lactate assay Kit has been used for the calibration curve. b, Michaelis-Menten curves of LDH at different pyruvate concentrations (0.1 to 1.3 mM) using 3.3 $\mu$M LDH as concentration in the droplets and supernatant (black), and 2 nM of free LDH in both supernatant phase (red) and buffer (cyan) in presence of 2 mM NADH. Inset of panel b: kinetic parameters values of LDH in the supernatant only. c, linear trends of lactate production over time at different pyruvate concentration (0.1-10 mM) of 3.3 $\mu$M LDH partitioned in the droplets. Volume fraction ($\phi$) used is 0.001 %.

ware (Mestrelab Research). The peaks at 3.02 ppm and 2.86 ppm stem from the two methyl groups in DMF; their integrals represent the signal from 6 protons per DMF molecule. The PEG peak appears at 3.71 ppm and its integral represents the contribution from approximately 354 protons per PEG molecule. The PEG concentrations were thereby determined from the relative height of the 3.71 ppm peak to the two DMF peaks at 3.02 ppm and 2.86 ppm and the known DMF concentration in the sample. Raw NMR data is included as supplementary data.

### Volume fraction

The volume fraction of the droplet phase at the chosen working conditions was determined using two independent methods. In the first method, we applied the lever rule to the previously measured BSA and PEG compositions of the droplet and supernatant phases, as well as the average composition of the droplet suspension. Specifically, the lever rule gives the droplet volume fraction, $\phi$, as

$$\phi = \frac{c_{a,i} - c_{S,i}}{c_{D,i} - c_{S,i}}, \tag{1}$$

where $c_{D,i}$ and $c_{S,i}$ are the mass concentrations of component $i$ in the droplet and the supernatant phases, respectively, $c_{o,i}$ is the average concentration of component $i$ in the droplet suspension, and $i \in \{BSA, PEG\}$. Using this formula, the droplet volume fraction can be calculated either from BSA or PEG concentration measurements. Because of the vicinity to the binodal line, we found that small variations of the average BSA concentration had a strong influence on the droplets' volume fraction, yielding a range from ($1\% < \phi_d < 5\%$). For our calculations, we used the average value $\phi = 3\%$.

In the second method, we prepared several mL of the droplet suspension, and separated the two phases by cen-

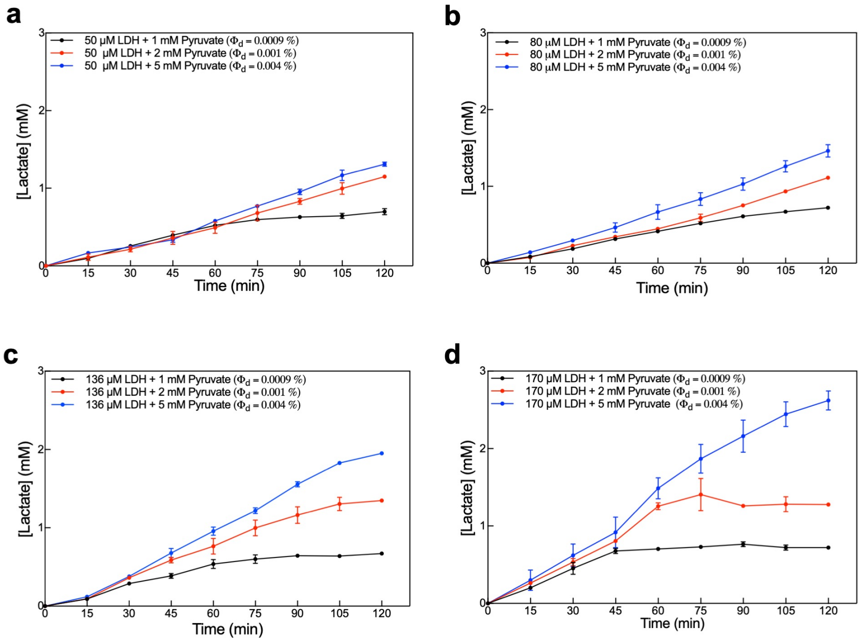


FIG. S4. **Supplementary Figure: Sustained Activity of LDH at different concentrations over time.** a, b, c ,d Sustained high metabolic activity adding 1.5, 2.4, 4.1 and 5.1 $\mu$M LDH respectively (50, 80, 136 and 170 $\mu$M inside the droplets), in presence of 10 mM NADH using different pyruvate concentration (1, black; 2, blue; and 5 mM red). Volume fraction ($\phi$) used in the experiments is reported in the legend of each panel.

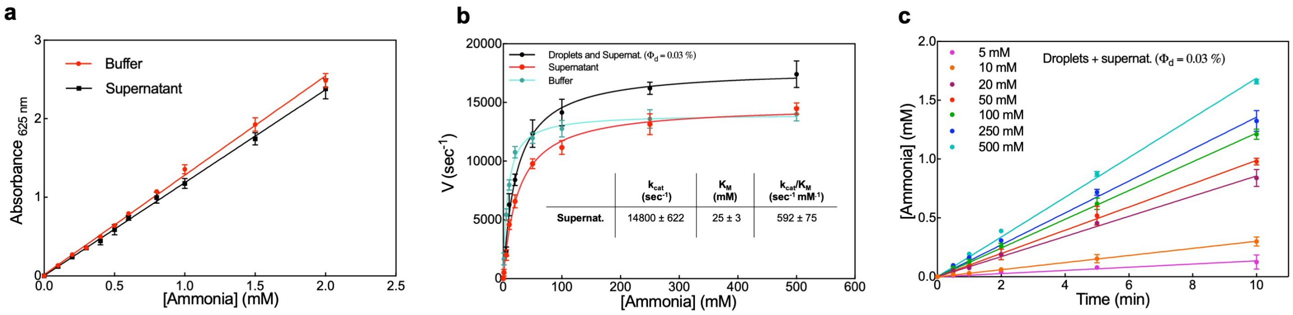


FIG. S5. **Supplementary Figure: Michaelis-Menten kinetics of urease in the droplets and supernatant, only supernatant and buffer.** a, Ammonia calibration curve in buffer (red) and in the supernatant (black) using $NH_4Cl$ solution from the Urease activity Assay Kit. b, Michaelis-Menten curves of urease at 1 $\mu$M concentration in droplets dispersed in supernatant (black), supernatant only (red) and in buffer, both at the final concentration of 0.3 nM. c, Activity of urease at different time points and urea concentrations ($5 - 500$ mM) using droplets containing 1 $\mu$M urease.

trifugation and directly compared the volumes of the two phases. We found $\phi$ values consistent with the lever rule.

### Sample chamber

In the presence of strong crowding by PEG, BSA droplets spread readily on bare glass and plastic substrates. Therefore, a surface coating is required to perform microscopy experiments. We coated microscopy glass slides with a thin poly(ethylene glycol) diacrylate (PEGDA) hydrogel layer [87]. Briefly, 24x55cm #1.5 thickness glass slides (VWR) were washed with water and ethanol three times, then cleaned with a UV-Ozone cleaner (ProCleaner, Bioforce Nanosciences) for a few minutes. The clean glass slides were pre-treated with a 0.3% v/v 3-(trimethoxysilyl)propylmethacrylate dispersed in a 5% v/v ethanol-water mixture. The solution was spread over the glass slides and the reaction allowed to proceed for 3 minutes, then it was quenched adding pure ethanol. The glass slides were stored in a dry box for 24h to allow the reaction to go to completion.

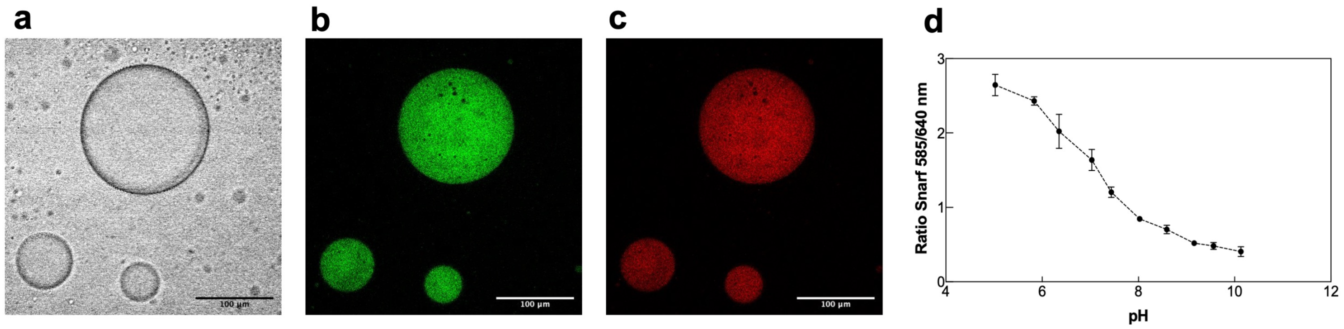


FIG. S6. **Supplementary Figure: Confocal imaging of BSA labeled with SNARF-1 in the droplets.** a, b, c, Confocal images of droplets containing BSA-SNARF-1 complex exciting the dye at $\lambda_g$ = 561 nm recording simultaneously the signal at $\lambda_g$ = 585 nm (green) and $\lambda_g$ = 640 nm (red). d, Calibration curve (ratiometric analysis) of BSA labelled with SNARF-1 inside the droplets at different pH (pH range 5.5-10.3) using potassium phosphate 0.1 M as buffer.

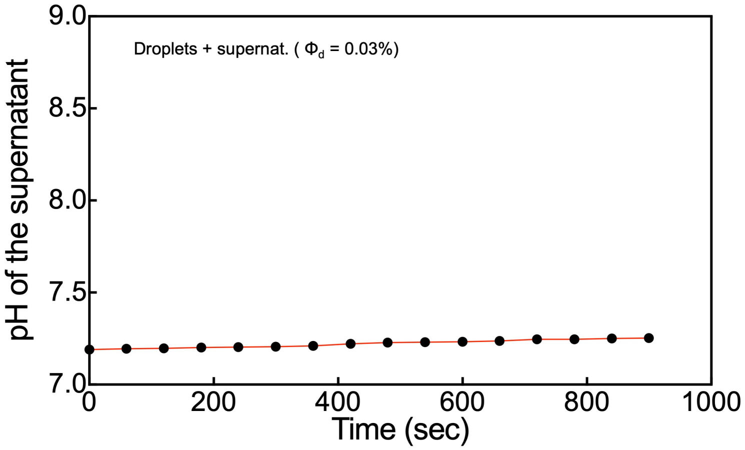


FIG. S7. **Supplementary Figure: pH of the droplets with partitioned urease diluted with the supernatant in presence of 100 mM urea.** Urease was used 1.0 $\mu$M in the droplets (the same experimental settings have been used during the measurements of the local pH change at the confocal microscope) and the pH of the solution has been measured using the pH meter (final volume was 180 $\mu$L, and $\phi = 0.03\%$).

We then prepared a solution at 1% w/w of the UV-activated initiator 2-Hydroxy-4'-(2-hydroxyyethoxy)-2-methylpropiophenone in water, and then added it to PEGDA 700 Da in a 80/20 initiator solution to PEGDA v/v ratio. 30 $\mu$L of the PEGDA-initiator mixture were pipetted on a pre-treated glass slide, and another clean, untreated glass slide (coated with an anti-rain film) was deposited on top. In this way, a thin layer of the PEGDA-initiator mixture was allowed to spread via capillary action between the two glass slides, and created a uniform coating. The glass slides were put under a UV lamp ($\lambda$ = 365 nm) for 20 minutes to allow complete curing of the PEGDA gel, then stored underwater. PEG-BSA droplets generally exhibit a contact angle slightly larger than 90 when resting on PEGDA-coated glass slides, as shown in Supplementary Fig. S15.

The well structure was 3D printed with a Prusa SL-1 3D printer (Prusa Research), with each well having 200 $\mu$L of final volume. Finally, the sample chamber was assembled bonding the PEGDA coated glass slide and the well structure with a UV-curable glue (Norland Optical Adhesive 63, Norland Products). The wells were sealed with Titer-Tops sealing film (Diversified Biotech) to avoid evaporation.

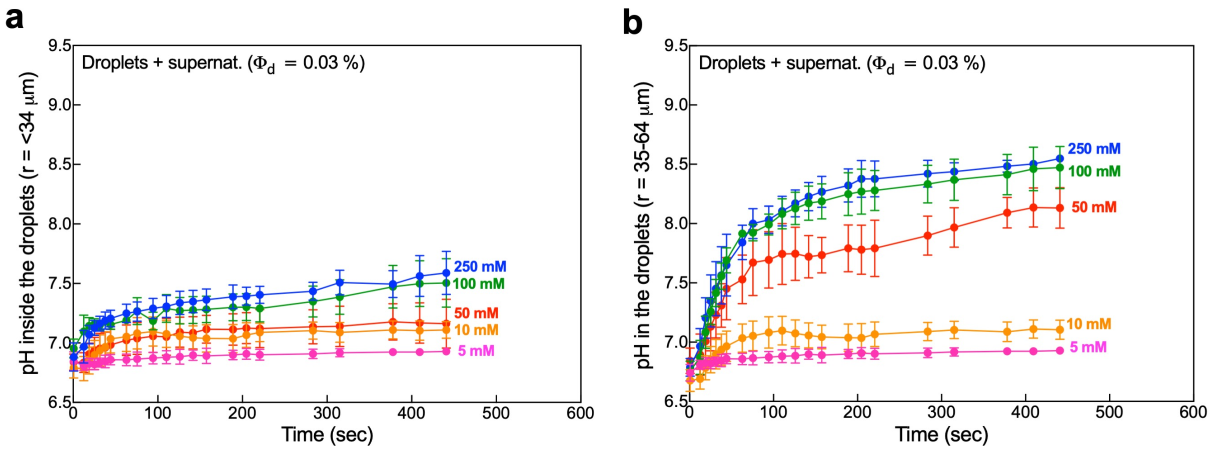


FIG. S8. **Supplementary Figure: pH change in the droplets versus time.** a, b, pH increase in the droplets of different size (a, $r \leq 34$ $\mu$m and b, between 35-64 $\mu$m) over time at different substrate concentration (5, 10, 50, 100 and 250 mM urea). The urease concentration in the droplets was 1.0 $\mu$M.

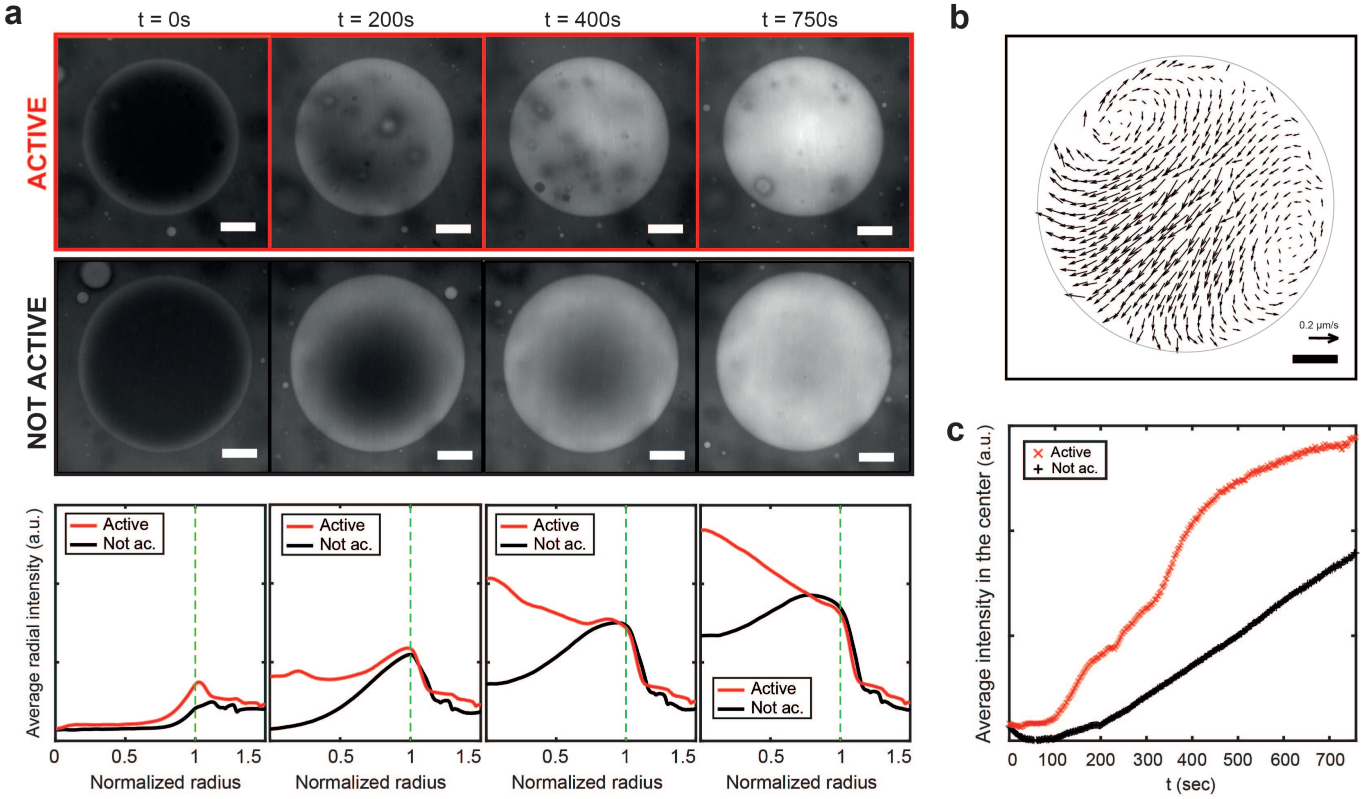


FIG. S9. **Supplementary Figure: Diffusion of rhodamine-B inside similarly sized urease active and non active droplets.** a. Top to bottom: fluorescent confocal images showing rhodamine distribution in an active droplet (urease concentration 1 $\mu$M, urea concentration 100mM) at different time frames versus a non active one, and corresponding radially averaged intensity profile plotted against the normalized radius. Green dotted line: droplet edge. b, Velocity field for the active droplet. c, Rhodamine intensity at the droplet center for the active and non active droplets, showing higher values for the active case. All the scale bars are 20 $\mu$m.

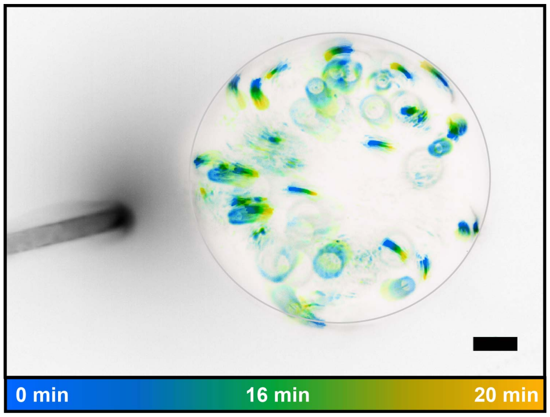


FIG. S10. **Supplementary Figure: Control experiment with micropipette.** Time projections of fluorescent intensity for a droplet and micro-pipette. The tracks show the color-coded trajectories of fluorescent tracers, while the dark halo the released solution. Scale bar: 20 $\mu$m.

#### Protein labelling

L-lactate dehydrogenase and Jack bean urease were tagged using Alexa Fluor 594 and 488 Microscale Protein Labeling Kit respectively, using the manufacturer's protocol but repeating the dialysis step three times to remove the free dye in the protein solution. The reaction between SNARF-1 and BSA was carried out in potassium phosphate 0.1 M pH 7.0 in presence of sodium bicarbonate 0.1 M for at least 1 hour at room temperature. Tagged proteins were further purified using membrane dialysis to reduce the free probe in solution. To determine the protein concentration of LDH and urease we used $\varepsilon_{280nm}$ = 43680 $M^{-1}cm^{-1}$ and $\varepsilon_{280nm} = 54165$ $M^{-1}cm^{-1}$, respectively (https://web.expasy.org/protparam/).

#### Droplet size distribution

We evaluated the size distribution of BSA droplets analyzing confocal microscopy z-stacks. Firstly, we made a droplet suspension at the standard working conditions, adding fluorescently labelled LDH at 30 $\mu$M concentration inside the droplets to allow their imaging in fluorescence. We then prepared a sample chamber sticking an imaging spacer (Secure-Seal, Grace Bio-Labs), diameter 9 mm, thickness 0.12 mm, to a coated glass slide. The thinner sample chamber was necessary to allow imaging of the sample chamber in the full z range. We then diluted the droplets 500-fold in substrate-free supernatant, and deposited 9 $\mu$L of the solution on the glass slide. The sample chamber was covered with a round glass slide top, and the edges additionally sealed with VALAP to limit evaporation.

Confocal microscopy z-stacks were acquired right after the sample preparation, using a 40X air objective, over a z range of 100 $\mu$m from the bottom of the sample chamber. Images were analyzed with Matlab, identifying for each droplet in the field of view the middle plane in 3D, and computing then the radial intensity average at that plane. The droplet radius was defined as the intensity width at half maximum of the radial intensity curve. In Supplementary Fig. S16 are reported the size distribution and the z-projection of a single field of view.

#### Kinetic parameters of LDH and activity measurements

The kinetic parameters ($k_{cat}$ and $K_m$ values) of LDH at the final concentration of 3.3 $\mu$M in the droplets and supernatant were assayed in presence of 2 mM NADH at different pyruvate concentrations (0.1 – 10 mM) over time, Supplementary Fig. S3c The $\phi$ value used for the determination of kinetic parameters of LDH was 0.001%. The droplets and supernatant has been diluted 1612-fold (0.62 $\mu$L) using the supernatant phase in a total volume of 1 ml at every substrate concentration. The $\phi$ value was achieved by diluting 0.62$\mu$L droplets phase into 1 mL supernatant phase at the appropriate substrate concentration. At each time point, the reaction mix was stopped using trichloroacetic acid (TCA) (10% final, 18 $\mu$L reaction mix + 2 $\mu$L of TCA 100%) and stored in ice. The stopped samples were then centrifuged for 10 minutes at 16900 g at room temperature and the amount of lactate in the reaction mix was determined using the Lactate Assay Kit (colorimetric mode) using a Multiskan Go plate reader (Thermo Fisher Scientific). Specifically, we ran the colorimetric lactate detection reaction as follows. First, we transferred 2 to 10 $\mu$L of the stopped samples to a 96-well plate and added Lactate Assay Buffer provided by the Lactate Assay kit to a final volume of 50 $\mu$L. Then, 50 $\mu$L of Master Reaction Mix (Lactate Probe + Lactate Enzyme Mix + Lactate Assay Buffer) were

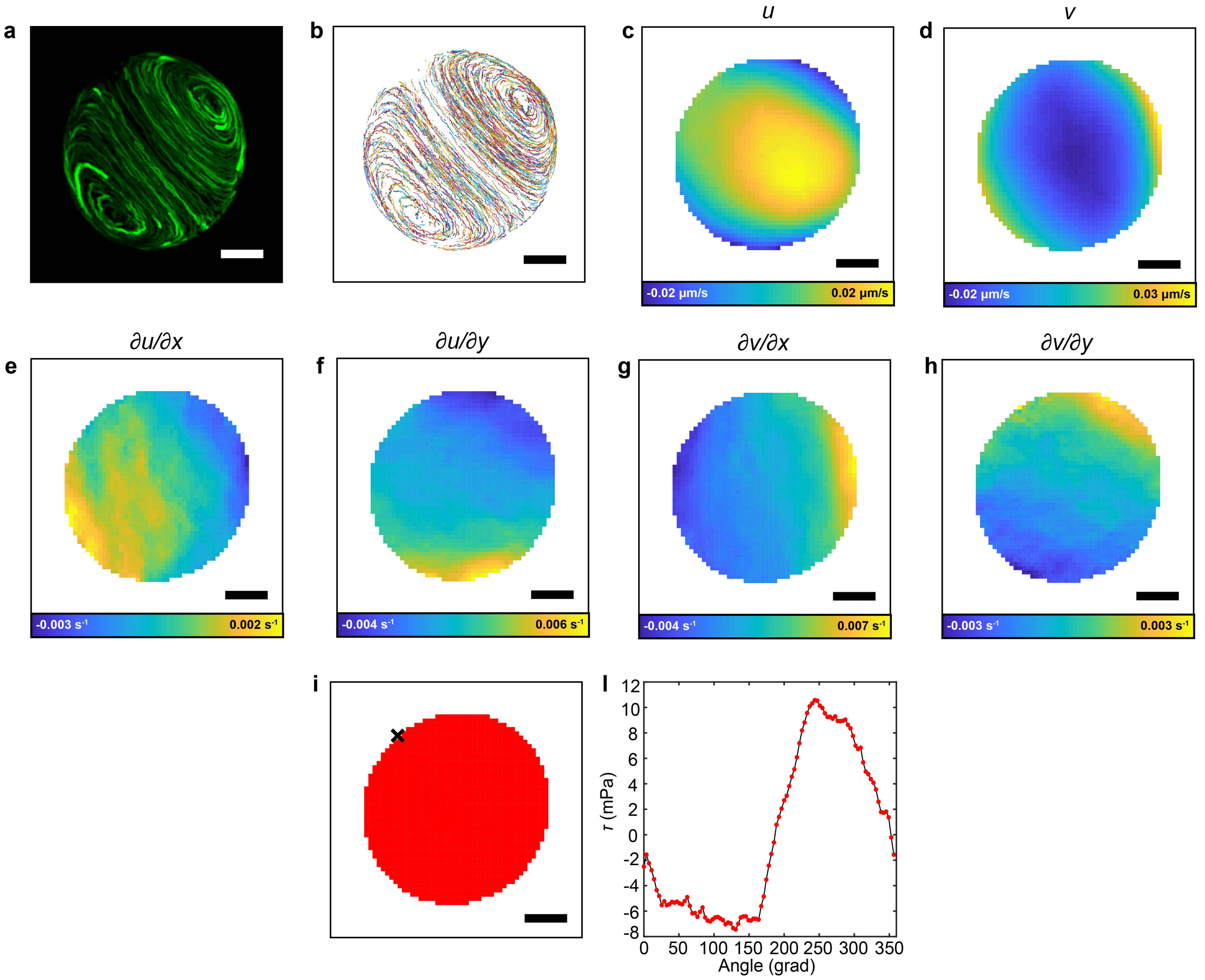


FIG. S11. **Supplementary Figure: Two-dimensional analysis of the flow and derivation of shear stress at $d = 3$ $\mu$m.** a, Time projections of fluorescent tracers over a period of 10 minutes and b, corresponding raw tracks. c,d, $x-$ and $y-$velocity fields, $u, v$. e-h, calculated velocity gradient fields. i, The black x represents the starting point for the calculation of the shear stress, moving counterclockwise along the droplet edge. l, Shear stress $\tau$ values along the droplets' edge. Scale bar: 10 $\mu$m

added, followed by shaking for 30 seconds. The reaction was subsequently incubated for at least 30 minutes while protecting the plate from the light and monitoring the absorbance at 570 nm every 5 minutes for an hour. Lactate solution provided by the Lactate Assay Kit was used to make the calibration curve in buffer and supernatant, as shown in Supplementary Fig. S3a ($0 - 10$ nmoles lactate is the range used for the calibration curve). The values of absorbance of the samples have been converted into velocity values for every pyruvate concentration and fitted (excluding the velocity values with substrate inhibition) using the Michaelis-Menten equation [30] as shown in Supplementary Fig. S3b. The $K_i$ values (substrate inhibition constants) were calculated by plotting the velocity values ($nmol_{\text{product}}/(\text{time} \cdot nmol_{\text{enzyme}})$) obtained at all pyruvate concentrations ($0.1 - 10$ mM) using the substrate inhibition equation [29].

To evaluate the partitioning of LDH, we measured the activity of the enzyme in the supernatant phase that remained after droplets were prepared and subsequently removed by centrifugation. Briefly, 1 mL of droplets suspension containing 60 nM LDH was split into two microcentrifuge tubes (500 $\mu$L each), and one tube was centrifuged for 30 minutes at 16900 g at room temperature. After transferring the supernatant to a new microcentrifuge tube, another centrifugation cycle was applied with the same settings as before. Then, 50 $\mu$L of supernatant containing final concentrations of 2 mM NADH and 1 mM pyruvate were added to 450 $\mu$L of droplets and supernatant, and supernatant alone. From this point

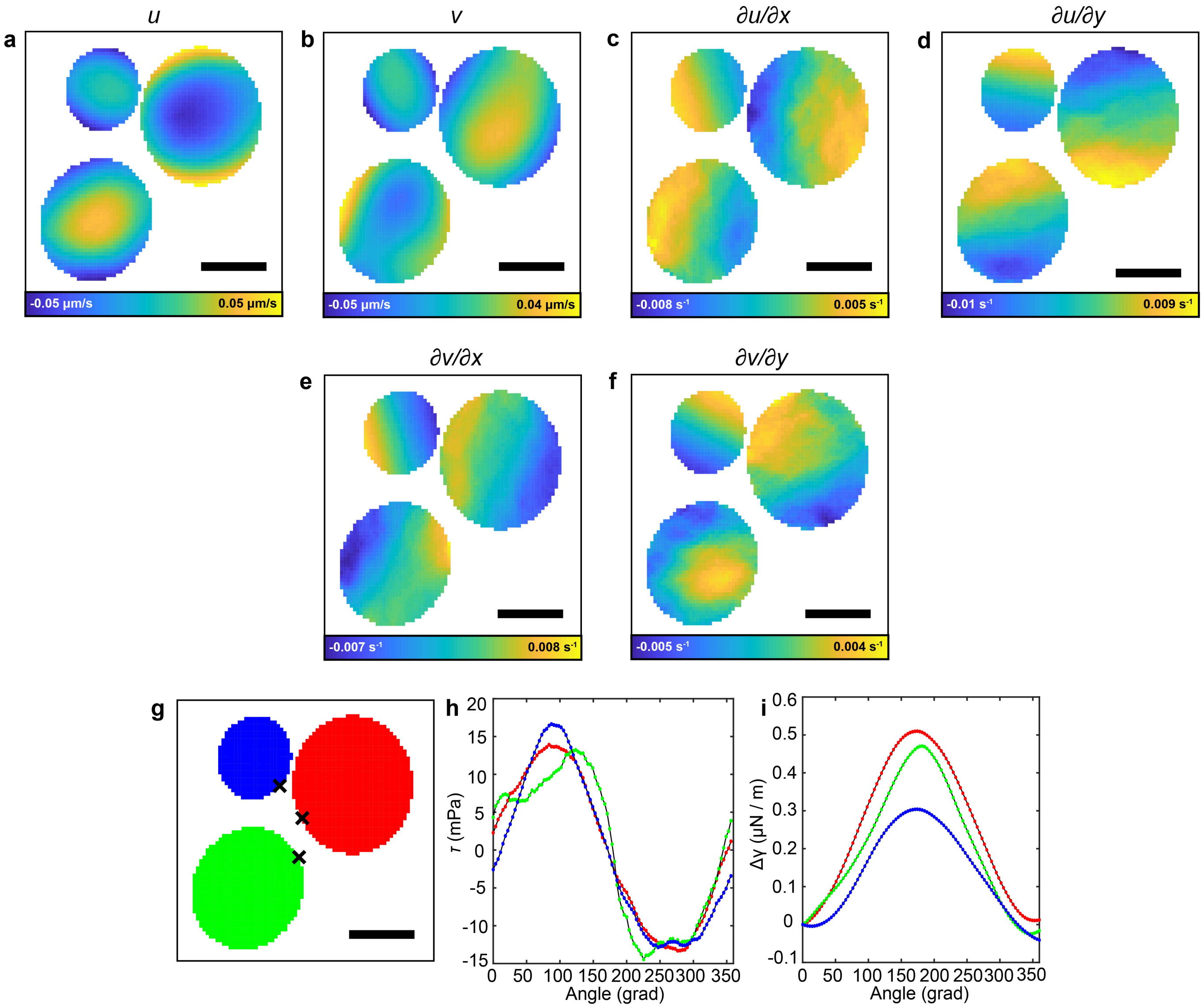


FIG. S12. **Supplementary Figure: Two-dimensional analysis of the flow and derivation of shear stress, surface tension for a group of three droplets, at $d = 4$ $\mu$m.** a, Velocity field in the x direction $u$. b, Velocity field in the y direction $v$. c, Derivative along the x axis of the velocity field in the x direction, $\partial u/\partial x$. d, Derivative along the y axis of the velocity field in the x direction, $\partial u/\partial y$. e, Derivative along the x axis of the velocity field in the y direction, $\partial v/\partial x$. f, Derivative along the y axis of the velocity field in the y direction, $\partial v/\partial y$. g, Color coding of the three droplets for the h, i plots. The black x represents the starting point for the calculation of the shear stress, moving counterclockwise along the droplet edge. h, Shear stress $\tau$ values along the droplets' edge. i, Delta surface tension $\Delta\gamma$ values along the droplets' edge. All scale bars are 20 $\mu$m.

on, we stopped, assayed and analyzed all the samples as previously described in this section. The buffer used for the activity measurements was potassium phosphate 0.1 M pH 7.0.

#### Metabolic rate: calculation and measurement

The specific metabolic rate expresses the amount of energy per unit time, per unit volume generated by metabolic reactions. We linked this quantity to enzymatic parameters through the reaction rate $v$ and reaction enthalpy $\Delta H$:

$$\dot{q} = v \cdot \Delta H \qquad (2)$$

The value of $\Delta H$ for the reaction of pyruvate conversion to lactate by LDH was 44 kJ/mol [88]. Considering that in most of our experimental conditions the substrate concentration is much larger than $K_m$, the reaction rate can be calculated from the estimated concentration of enzyme in the droplets $[E]_d$ and the enzyme turnover

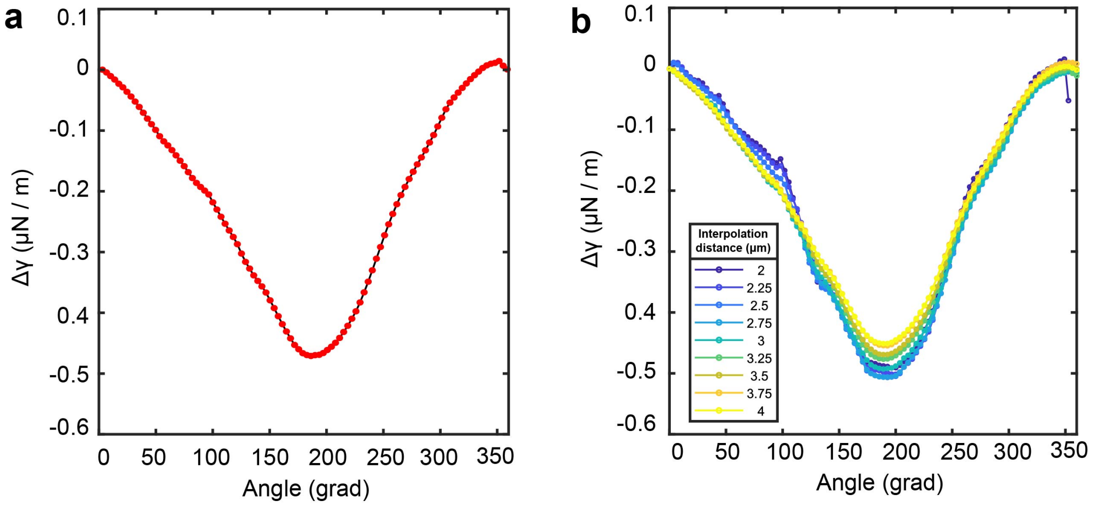


FIG. S13. **Supplementary Figure: Measuring surface tension differences around droplets.** a, Surface tension difference calculated around the droplet in Figure S11, using $d = 3$ $\mu$m. b, Surface tension differences for the same droplet, calculated using values of $d$ ranging from 2 to 4 $\mu$m. The results are insensitive to the value of $d$ in this range.

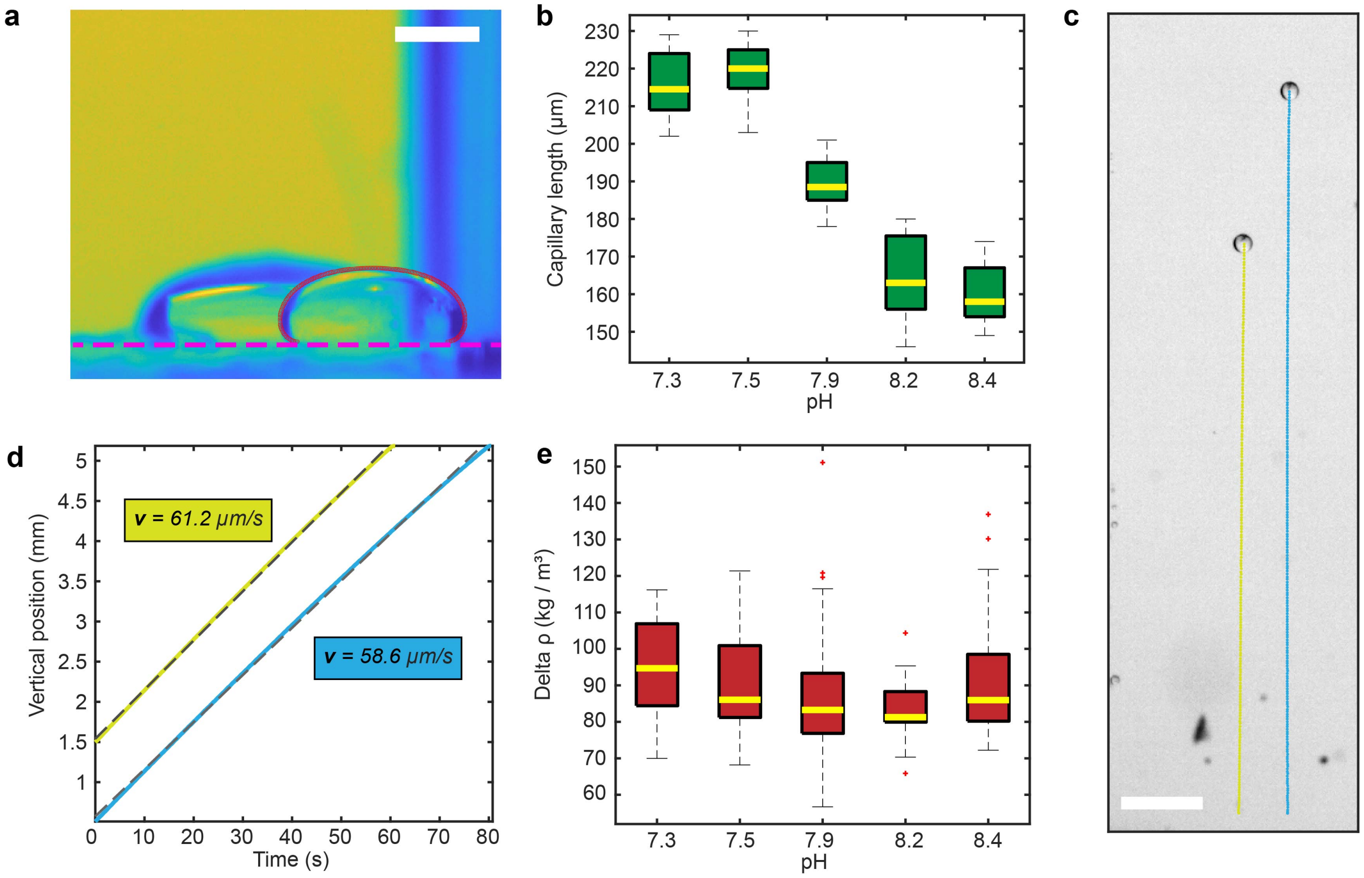


FIG. S14. **Supplementary Figure: Measurement of equilibrium surface tension.** a, Brightfield image of a droplet from the side with fitted profile (in red). Purple dotted line: glass slide surface. Scale-bar: 500 $\mu$m b, Box plot of the calculated capillary length versus pH of the supernatant. For each point 20 droplets were analyzed. c, Trajectories of two falling droplets as resulting from tracking. Scale-bar: 500 $\mu$m d, Blue and yellow lines: tracked vertical position versus time of the two droplets of panel c. Dashed gray line: linear fit. The limit velocities have been extrapolated from the curves slope. f, Density difference between the supernatant and the droplet phase calculated from the terminal velocity. For each point 20 droplets were analyzed.

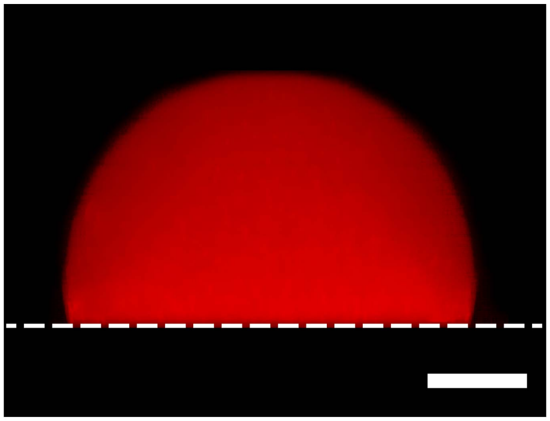

FIG. S15. **Supplementary Figure: Side view of a droplet resting on a PEGDA-coated glass slide.** Confocal view of a Rhodamine-B containing droplet resting on a PEGDA-coated glass slide. Dashed line: glass slide top surface. Scale bar: 15 $\mu$m.

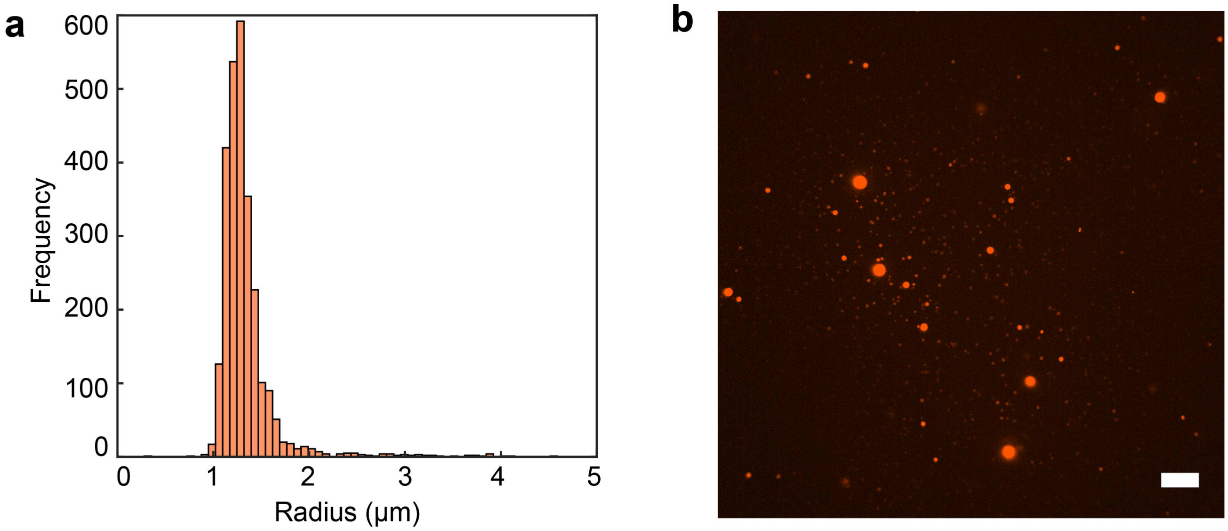


FIG. S16. **Supplementary Figure: Size distribution of BSA droplets.** a, Histogram of the radii distribution of BSA droplets diluted 500 times in supernatant. b, z-projection of confocal z-stack for BSA droplets diluted 500 times in supernatant. Scale bar: 25 $\mu$m.

number $k_{cat}$ as follows:

$$v = [E]_d \cdot k_{cat} \quad (3)$$

The measurement of enzymatic activity shows that enzymes used in this work partition remarkably well in the droplet phase (Fig 1c, 2b): hence, for our calculations we can assume perfect partitioning. The enzyme concentration in the droplets was therefore estimated from the known enzyme concentration in solution $[E]_s$ and the droplet volume fraction $\phi_d$ as follows:

$$[E]_d = \frac{[E]_s}{\phi_d} \quad (4)$$

Considering the volume fraction measured in the standard conditions is $\phi_d = 3\%$ (see previous sections), $[E]_d$ is roughly 33 times $[E]_s$ in our experiments. The concentration of enzyme in the droplets throughout the present work was calculated in this way.

The turnover number $k_{cat}$ for these experiments was estimated, with the known enzyme concentration in the droplets, from the slope of the lactate production curves versus time in linear regime at timepoint $t'$ (Supplementary Fig. S4) as follows:

$$k_{cat} = \frac{[LAC]_{t'}}{t' \cdot [E]_d} \quad (5)$$

In particular, we used $t' = 45$ minutes as it is the latest timepoint (i.e. with the highest measurable signal) in which all of the curves are in linear regime. At times later than 45 minutes some combinations of enzyme and substrate concentration show curve flattening due to substrate consumption.

Experimentally, the droplets were prepared using different enzyme concentrations $[E]_s = 1.5$, 2.4, 4.1 and 5.1 $\mu$M. The corresponding $[E]_d$ were calculated to be roughly 50, 80, 136 and 170 $\mu$M. The droplets were gently mixed in 1 mL of supernatant containing 10 mM NADH and different pyruvate concentrations (1, 2 and 5 mM). The reactions were stopped and the lactate concentration quantified using the Lactate Assay Kit as described in the previous section (Supplementary Fig. S4).

#### Activity measurements and kinetic parameters of urease

0.2 mL of droplets solution containing 1 $\mu$M (partitioned) urease were prepared and divided equally (100 $\mu$L) into two different microcentrifuge tubes. One tube was centrifuged twice (as described in the previous section) and then 1 $\mu$L of the recovered supernatant was tested for the enzymatic activity by adding 99 $\mu$L of supernatant containing 100 mM urea. The same procedure (without the centrifugation step) was employed for the solution containing the droplets. At each time point 100 $\mu$L of reaction mix was stopped by adding 500 $\mu$L of 5 % of phenol nitroprusside solution immediately followed by 500 $\mu$L of 2 % alkaline hypochlorite solution. The mixture was then incubated for at least 20-30 minutes at room temperature, thereby allowing the colored complex to develop. The ammonia generated by each reaction was then determined by measuring absorption at 625 nm using a Multiskan Go plate reader and compared to a standard curve made with $NH_4Cl$ of known concentrations (Supplementary Fig. S5a). The same procedure was employed for the determination of the kinetic parameters of urease in droplets (enzyme concentration in the droplets 1 $\mu$M, Supplementary Fig. S5c), in supernatant only, and in buffer using different substrate concentrations (1 - 500 mM). The kinetic parameters of urease ($k_{cat}$ and $K_m$ values) were calculated by plotting the velocity values ($nmol_{\text{product}}/(\text{time} \cdot nmol_{\text{enzyme}})$) using the Michelis-Menten equation [30] as reported in Supplementary Fig. S5b.

#### pH Imaging

We quantified *in situ* pH variations using fluorescence ratiometric imaging. After labelling BSA with SNARF (as described above), it was used to make droplets following the usual procedure, with a concentration of 1 $\mu$M urease in the droplets. Because the labelling degree of BSA was low (due to the the high DMSO concentration used during SNARF resuspension and the related side effects on proteins using high DMSO concentrations [89, 90], we added to 0.5 mL of SNARF-1-labelled BSA droplets ($\phi = 0.03$) additional SNARF solution at a final concentration of 50 $\mu$M. The droplets solution was then centrifuged for 30 seconds at 15000 rpm at room temperature and roughly half of the supernatant was removed from the microcentrifuge tube. The droplets phase was gently resuspended and 1-2 $\mu$L was diluted in 9-18 $\mu$L of substrate and BSA-SNARF free supernatant in a microcentrifuge tube. A confocal microscope LSM 880 Airyscan (Carl Zeiss Microscopy) has been used to image the pH inside the droplets. Depending on the size of the droplets, two different objectives have been used: a 40x/1.30 PlanApo oil lens and a 63x/1.45 PlanApo oil for larger droplets and smaller ones, respectively. Fluorescence imaging was performed exciting the dye at 561 nm and monitoring the pH-dependent emission spectral shifts simultaneously in two separate channels. The emission wavelengths selected were $\lambda_g = 585$ nm and $\lambda_r = 640$ nm.

For the pH measurements of Fig. 2, 10-20 $\mu$L of diluted droplets were loaded into the sample chamber and the enzymatic reaction was started adding 150 $\mu$L of supernatant containing the urea at different concentrations. For calibration purposes, droplets were prepared and loaded into the chamber as above, but we added 150 $\mu$L of pH adjusted supernatant without urea. The droplets were allowed to incubate for one hour before imaging. The resulting calibration curve is shown in Supplementary Fig. S6).

#### Particle tracking microrheology

In order to measure the viscosity of the droplet phase, green fluorescent polystyrene nanoparticles tracers of 0.2 $\mu$m average diameter were added to the droplet suspension, at final concentration of 0.0005% solid content. The nanoparticles displayed excellent partitioning in the droplet phase, with only minimal aggregation visible (Supplementary Fig. S1a). We took timelapses of fluorescent tracers at the confocal microscope (time interval = 2 s) and tracked their motion in 2D (Supplementary Fig. S1b). The mean squared displacement (MSD) of the particles increased linearly in time, implying that the droplets behave as simple Netwonian liquids. The diffusion coefficient was determined from the slope of MSD, and the viscosity was determined using the Stokes-Einstein relation:

$$\eta = \frac{k_B T}{6\pi D r} \tag{6}$$

With $k_B$ the Boltzmann constant, $T$ the absolute temperature and $r$ the radius of the particles.

Firstly, we performed the measurement of the baseline viscosity for BSA droplets at the chosen working conditions (without presence of enzymes, co-factors or substrates). In Supplementary Fig. S1c we report the MSD curves for several different droplets. The different curves overlap quite well in the interval 10 to 100 seconds.

In Supplementary Fig. S1d we also present the diffusion coefficient kernel probability distribution for each experiment. The averaged ensemble diffusion coefficient for the droplets is $D = 5.3 \cdot 10^{-4} \pm 0.4 \cdot 10^{-4}$ $\mu$m$^2$/s, with corresponding viscosity value $\eta = 2.1 \pm 0.2$ Pa s, roughly 2000 times the viscosity of water. Thus, BSA droplets are highly viscous domains characterized by purely viscous behavior at this length scale.

We also evaluated the viscosity of droplets with partitioned LDH, at a concentration of roughly 80 $\mu$M in the droplets. In Supplementary Fig. S1e,f we report the MSD curves for several different and the diffusion coefficient kernel distribution for the different droplets. The averaged ensemble diffusion coefficient is $D = 4.9 \cdot 10^{-4} \pm 0.5 \cdot 10^{-4}$ $\mu$m$^2$/s, with corresponding viscosity value $\eta = 2.2 \pm 0.2$ Pa s. Thus, the viscosity values of droplets with and without LDH are virtually identical.

Finally, we have assessed the viscosity of the supernatant. As most of the macromolecular components are partitioned into the droplets, we do not expect any significant variation of its viscosity between different experiments with other components present. For this experiment, we have isolated the supernatant of a sample at the chosen working conditions (without presence of enzymes, co-factors or substrates) as previously described. We then added fluorescent polystyrene nanoparticles of 1 $\mu$m diameter with a final solid fraction of 0.0001%. We then acquired timelapses of particles diffusion (time interval = 300 ms) and analyzed them as previously described. In Supplementary Fig. S1h we report the MSD of several different experiments. All curves are linear, and the calculated ensemble diffusion coefficient it $D = 0.017 \pm 0.001$ $\mu$m$^2$/s, with corresponding viscosity value $\eta = 0.012 \pm 0.006$ Pa s, roughly 10 times the viscosity of water. This value agrees with reported measurements of viscosity for PEG solutions at similar molecular weights and concentrations [91].

Concluding, the viscosity of the droplets is roughly two orders of magnitude larger than the one of the supernatant, as their ratio is:

$$\frac{\eta_d}{\eta_s} \approx \frac{2Pa \cdot s}{0.012Pa \cdot s} \approx 167 \tag{7}$$

### Rhodamine-B uptake

In order to test the rhodamine-B uptake in the droplets we made a 1 mg/mL rhodamine-B solution in Milli-Q water. We then prepared a few milliliters of supernatant containing urea as previously explained, and added the rhodamine solution to a final concentration of 0.01 mg/mL. Later, two different droplets suspension were made, one with $[E]_s = 0.03$ $\mu$M urease, ($\approx 1$ $\mu$M in the droplets) for the active case, and one without enzyme for the non-active case. Both droplets suspensions also contained fluorescent nanoparticles (200 nm, see Supplementary Materials ) at a final concentration of 0.0005% solid fraction. The droplets were re-dispersed in substrate and rhodamine free supernatant and pipetted in the sample chamber. Then, the rhodamine-B containing supernatant was added. The volume ratio between supernatants and the urea concentration was set to have a final 100 mM concentration of urea.

Confocal microscopy time-lapses were acquired in both the rhodamine and tracers channels (time delay = 2 sec) to assess both the dye diffusion (Supplementary Fig. S9a) and the fluid flow (Supplementary Fig. S9b). To directly compare the active and non-active case we performed experiments on a high number of droplets and then matched the ones with similar diameter.

As evident from Supplementary Fig. S9a, advection visibly alters the rhodamine-B distribution. This is visible both from confocal microscopy images, and from the calculated radial average of the rhodamine fluorescent intensity in the droplets. This observation is additionally confirmed by the evaluation of the average intensity calculated in the center of both droplets (Supplementary Fig. S9c). The maximum difference in intensity is reached at around 400 seconds, where the active droplets shows more then double the value of the inactive one.

We estimated the diffusion coefficient of rhodamine-B inside the droplets from the time-evaluation of its intensity profile. From the data in Supplemental Fig. (S9a), one can see that it takes about 200 seconds for the rhodamine intensity half-way to the center of the droplet (about 30 $\mu$m) to reach about half its maximum value. From the relationship between mean square displacement (MSD) and diffusion coefficient,

$$\mathrm{MSD} = 2Dt \tag{8}$$

one can estimate the diffusion coefficient to be about $D \approx 2$ $\mu$m$^2$/s, roughly 200× lower than the reported value for water [92].

### Micro-pipette experiments

Solutions for micro-pipette experiments was prepared starting from a KP solution at pH = 10. First, we prepared a KP solution at 500 mM and pH 7.5 (at the highest end of the KP buffering window). Then we further shifted the pH to alkaline adding NaOH 1M solution drop-wise until a value of 10 was reached.

Afterwards, we prepared a droplets suspension using the alkaline buffer in place of the one at neutral pH and isolated the supernatant, as previously reported. In order to make the gradient visible with fluorescence microscopy, we prepared a 1 mg/mL FITC solution in dimethyl sulfoxide (DMSO), and added it to the supernatant at a final concentration of 0.005 mg/mL. The final pH of the supernatant measured with a pH meter was around 8.4.

Then, we prepared a urea-free droplet suspension, adding fluorescent nanoparticle tracers (diameter 0.2 $\mu$m, final concentration 0.0005% solid content) to visualize the inner flow, and placed it into a sample chamber with an open side. We loaded the alkaline buffered solution into a glass micro-pipette with a tip of roughly 10 $\mu$m of outer diameter and inserted it in the sample chamber, locating it in the vicinity of a droplet. With the aid of a micro-pressure controller, we then gently released the alkaline solution and recorded 20 minutes timelapses of the motion at a fluorescence microscope. We estimated the flow rate in the pipette based on its inner diameter ($\approx 5$ $\mu$m) and the velocity of the fluorescent fluid inside the pipette to be around 10 $\mu$m$^3$/s.

As a control experiment, we prepared an identical solution as described above but using KP at pH 7 instead of 10. Both the final pH of the supernatant and of the droplet suspension was measured to be 7.2. Upon release of the solution, a top flow rate of about 0.01 $\mu$m/s was observed inside the droplets, about a quarter of the speed observed in the presence of the pH gradient (Supplementary Fig. S10). Notably, its direction was *opposite* to the one recorded for the alkaline buffered solution.

#### Measurement of shear stresses at droplet surfaces

We measure shear stresses normal to the droplet surface, $\tau^{\text{in}}$, by measuring the local strain rate, $\epsilon^{\text{in}}$, and relating them via

$$\tau^{\text{in}} = \mathbf{t} \cdot \sigma^{\text{in}} \cdot \mathbf{n} = 2\eta^{\text{in}} \mathbf{t} \cdot \epsilon^{\text{in}} \cdot \mathbf{n}. \quad (9)$$

Here, $\mathbf{t}(x, y) = (t_x, t_y)$ is the local tangent vector to the droplet surface, and $\mathbf{n}(x, y) = (n_x, n_y)$ is the normal vector pointing out of the droplet. $\eta$ is the viscosity, $\sigma$ is the stress tensor, and the strain rate is $\epsilon = \nabla \mathbf{u} + \nabla \mathbf{u}^T$, with $\mathbf{u}(x, y) = (u, v)$ being the local velocity field in Cartesian coordinates. The superscript 'in' indicates quantities just inside the droplet surface. In terms of velocity gradients, this becomes

$$\tau^{\text{in}} = 2\eta^{\text{in}}(t_x, t_y) \cdot \begin{pmatrix} \frac{\partial u}{\partial x} & \frac{1}{2}\left(\frac{\partial u}{\partial y} + \frac{\partial v}{\partial x}\right) \\ \frac{1}{2}\left(\frac{\partial u}{\partial y} + \frac{\partial v}{\partial x}\right) & \frac{\partial v}{\partial y} \end{pmatrix} \cdot \begin{pmatrix} n_x \\ n_y \end{pmatrix}. \quad (10)$$

To evaluate this expression, we use the tracks of the fluorescent tracer particles in droplets (Supplementary Fig. S11a, b). We assume that the flow is steady, so that velocity data from each particle at every time point can be used to create a velocity map inside of the droplets. We then interpolate this data at regularly spaced grid points around the edge of droplets to give local velocities (Supplementary Fig. S11c, d) and velocity gradients (Supplementary Fig. S11e-h). We perform this interpolation by taking all the particle tracks within a distance $d$ of each grid point, and performing a linear interpolation of the particle velocity data. We also perform the interpolation at regular grid points inside of the droplet to visualize the flow fields. $d$ must be chosen carefully to be smaller than all the features in the flow, but large enough that there are sufficient particle tracks to do an accurate interpolation. We show below that the results are insensitive to the value of $d$ in a certain range. The final, measured shear stresses for this droplet are given in Supplementary Fig. S11l.

#### Dynamic Measurement of Surface Tension Differences

Shear stresses at the droplet interface can either be caused by Marangoni stresses, or by shear from a flow outside of the droplet. This is captured by the Marangoni equation representing stress balance at the interface:

$$\frac{d\gamma}{ds} = -\tau^{\text{out}} + \tau^{\text{in}} = -2\eta^{\text{out}} \mathbf{t} \cdot \epsilon^{\text{out}} \cdot \mathbf{n} + 2\eta^{\text{in}} \mathbf{t} \cdot \epsilon^{\text{in}} \cdot \mathbf{n}. \quad (11)$$

Here, $\gamma$ is the surface tension, $s$ is arclength around the droplet surface, and superscript 'out' corresponds to parameters in the liquid just outside of the droplet surface.

We can simplify this by noting that, as $\eta^{\text{in}}$ is much bigger than $\eta^{\text{out}}$ (as described in the previous sections), $\tau^{\text{in}} \gg \tau^{\text{out}}$, and so

$$\frac{d\gamma}{ds} = \tau^{\text{in}}. \quad (12)$$

Thus, we calculate surface tension differences, $\Delta\gamma(s)$, along the droplet surface by setting the calculated shear stresses inside the surface of the droplet to $d\gamma/ds$, and numerically integrating around the droplet (*e.g.* [93]). Supplementary Fig. S13b shows this for a range of values of $d$. We see that the results are essentially independent of $d$ between 2 and 4 $\mu$m. This range makes sense, as it is small in comparison to the bulk features of the droplet velocity fields (see Supplementary Fig. S11c,d).

There is one important check to ensure that the calculation is consistent with the presence of Marangoni stresses. As a material property, $\gamma$ has a unique value at each point on the surface. Equivalently, $\Delta\gamma(S) = 0$, where $S$ is the total arclength around the droplet perimeter. If this does not hold, there is either an error in the calculation, or there is missing physics in Eq. 12. Here, for all droplets, we find that $\Delta\gamma(S) \approx 0$ independent of the choice of $d$, giving us confidence in the calculation (see *e.g.* Supplementary Fig. S13a).

#### Equilibrium measurement of surface tension

As previously reported the surface tension at the droplet-supernatant interface can be calculated from two measurements [47]: by analyzing the shape of droplets resting on a glass slide (from which we derive the capillary length, $L_c$), and their sedimentation velocity (from which we derive the density difference between the two phases, $\Delta\rho$) and applying the following equation:

$$\gamma = L_c^2 \cdot \Delta\rho \cdot g \tag{13}$$

We prepared five KP solutions at progressively more alkaline pH (7, 7.5, 8, 9, 10) as previously described. We then used these buffers to prepare droplets suspensions free of substrates, enzyme or fluorescent tracers: only PEG, BSA, KCl and the KP were added, at the chosen concentrations. The final pH of the supernatant for these samples measured with a pH meter was 7.3, 7.5, 7.9, 8.2 and 8.4 respectively. We then centrifuged the tubes at 16900 g for 30 minutes, and separated the supernatant from the droplet phase.

Around 2 mL of the supernatant at the pH of choice were loaded in a polystyrene semi-micro cuvette, and the corresponding droplets were added from the top using a positive displacement pipette. Both the sedimented and falling droplets were imaged with the optical setup described in [47].

While the capillary length was derived from the droplet shape (Fig. 3f, Supplementary Fig. S14a, b), the density difference was calculated tracking the motion of falling drops in the supernatant-filled cuvette, and calculating the terminal velocity fitting the vertical change in position versus time (in the approximation case of $\eta_D \gg \eta_{SN}$, ), as reported in Supplementary Fig. S14c, d. The value of $\Delta\rho$ (91.6 kg/m$^3$) was found to be essentially independent from the pH conditions (Supplementary Fig. S14e), and well in the range of typical protein droplets aqueous suspensions [47].

#### Transport limitations

Here, we develop simple transport models to estimate the profiles of concentration and temperature of droplets hosting chemical reactions. We consider a spherical reactive domain of radius R (the droplet), surrounded by an unbounded domain (the supernatant) [32, 33]. The conversion of substrate, $S$, into product, $P$, catalyzed by enzyme, $E$, follows a two-step reaction via

$$S + E \underset{k_r}{\overset{k_f}{\rightleftharpoons}} ES \xrightarrow{k_{cat}} P + E, \tag{14}$$

where $ES$ denotes the enzyme-substrate complex. Under our experimental conditions, $c_S \gg c_E$ (free ligand approximation), where $c_S$ is the substrate concentration and $c_E$ is the enzyme concentration. Furthermore, $k_r \gg k_{cat}$ allows us to make the quasi steady-state approximation, *i.e.* $c_{ES} \approx$ const., where $c_{ES}$ is the concentration of the enzyme-substrate complex. We can then write down the net reaction rate, $s$, obeying Michaelis-Menten kinetics as

$$s = -\frac{dc_S}{dt} = k_{cat} c_E \frac{c_S}{K_M + c_S}, \tag{15}$$

where $k_{cat}$ is the catalytic rate constant, and $K_M = (k_r + k_{cat})/k_f$ is the Michaelis constant [30]. In the absence of advective flows, the transport of substrate, enzyme, and product is governed by diffusion via

$$\mathbf{j}_i = -D_i \nabla c_i, \tag{16}$$

where $i \in \{E, S, P\}$ and the off-diagonal (cross-diffusion) terms of the diffusion tensor are assumed to vanish. At steady-state, the concentration profile of the substrate is therefore well-approximated by

$$D_S \nabla^2 c_S = k_{cat} c_E \frac{c_S}{K_M + c_S}, \tag{17}$$

We first develop a simple analytical solution in the limit of high substrate concentrations, $c_S \gg K_M$. Here, the right hand side of Eq.17 simplifies to $k_{cat} c_E$. Assuming that $c_E \approx$ const. for $r < R$ and $c_E \approx 0$ for $r > R$, we can write down the solution for the steady-state substrate concentration as

$$c_S(r) = \begin{cases} c_S^\infty + \frac{1}{2}\frac{k_{cat} c_E R^2}{3 D_S^{in}} \left( \left(\frac{r}{R}\right)^2 - 1 - 2\frac{D_S^{in}}{D_S^{out}} \right), & \text{if } r < R \\ c_S^\infty - \frac{k_{cat} c_E R^2}{3 D_S^{out}} \left(\frac{R}{r}\right), & \text{if } r > R, \end{cases} \tag{18}$$

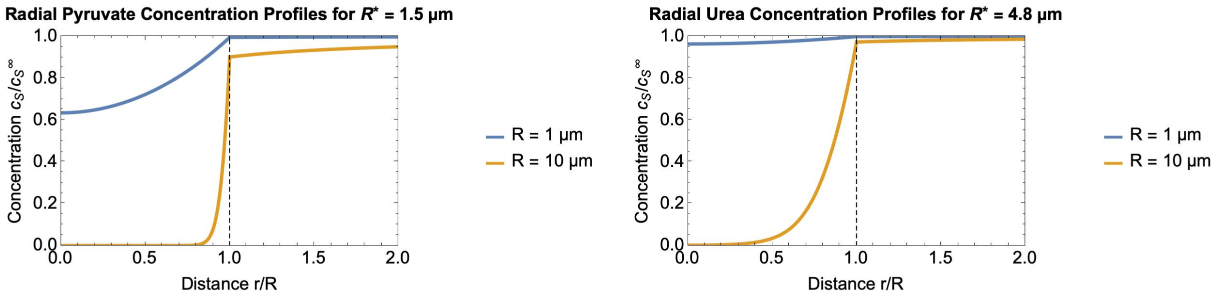


FIG. S17. **Supplementary Figure: Steady-state substrate concentration profiles.** Numerical solutions of Eq. 17 are plotted for $R = 1$ $\mu$m and $R = 10$ $\mu$m. *(left)* Calculation for pyruvate/LDH using $c_S^\infty = 1$ mM, $c_E = 3$ $\mu$M, $D_S^{out} = 83.5$ $\mu$m$^2 s^{-1}$, $D_S^{in} = 0.5$ $\mu$m$^2 s^{-1}$, $k_{cat} = 469$ s$^{-1}$, and $K_M = 0.22$ mM. *(right)* Calculation for urea/urease using $c_S^\infty = 100$ mM, $c_E = 1$ $\mu$M, $D_S^{out} = 115$ $\mu$m$^2 s^{-1}$, $D_S^{in} = 0.7$ $\mu$m$^2 s^{-1}$, $k_{cat} = 18 \times 10^3$ s$^{-1}$, and $K_M = 23$ mM.

where $D_S^{in}$ and $D_S^{out}$ are the diffusion constants of the substrate inside and outside the droplet, respectively, and $c_S^\infty$ is the substrate concentration in the supernatant. In order for the substrate concentration at the center of the droplet to be similar to the bulk, *i.e.*

$$c_S^\infty - c_S(r=0) \ll c_S^\infty, \tag{19}$$

the droplet size must be small. Specifically,

$$R \ll R^* = \sqrt{\frac{6 c_S^\infty D_S^{in}}{k_{cat} c_E}}. \tag{20}$$

Values of $R^*$ corresponding to our experimental conditions are reported in Fig. S17.

We now solve Eq. 17 numerically to reveal the expected concentration profiles in our experimental conditions. To do so, we need to specify diffusion coefficients of pyruvate and urea inside and outside the droplets. In the absence of direct measurements for these two quantities, we scale their published diffusion coefficients in water by the relative viscosity of the droplet and continuous phases, 2000× and 12×, respectively. For the diffusion coefficient of urea in water, we used $D_{\mathrm{urea,w}} = 1373$ $\mu$m$^2$s$^{-1}$ and for pyruvate we used $D_{\mathrm{pyruvate,w}} \approx 1000$ $\mu$m$^2$s$^{-1}$ [94, 95].

The solutions for urease and LDH are shown in Fig. S17. We have plotted the solutions for 1 $\mu$m and 10 $\mu$m radius droplets under our experimental conditions. For both urease and LDH, the reaction occurs throughout for the droplet only for the smaller size. Note that these solutions ignore substrate inhibition (observed for LDH) as well as advection (observed for urease droplets).

Finally, we estimate the temperature change due to heat released by the enzymatic reactions. Using similar assumptions as those used to derive Eq. 17, we can solve the heat equation and derive an upper estimate for the temperature increase at the center of droplets, $\Delta T^\circ$. Specifically, we find that

$$\Delta T^\circ \approx \frac{\dot{Q} R^2}{2\kappa}, \tag{21}$$

where $\kappa$ is the thermal conductivity [32, 33]. With $\dot{Q} = 10$ MWm$^{-3}$, $\kappa = 0.6$ WK$^{-1}$ for water and $R = 10$ $\mu$m, we estimate that $\Delta T^\circ \approx 0.8$ mK.

#### Theoretical Description of the Chemically Activated Flow

Here we provide the details of the theoretical model that we use to explain the origin of the flow and calibrate using the pipette experiment in which activity is induced within the droplet that encapsulates enzymes.

##### *Chemical Field*

We specifically consider a chemical source or sink with flux $I$, located at distance $h$ from the centre of a droplet of radius $R$, as shown in Fig. S18(a). The concentration field of the relevant chemical species is governed by the diffusion

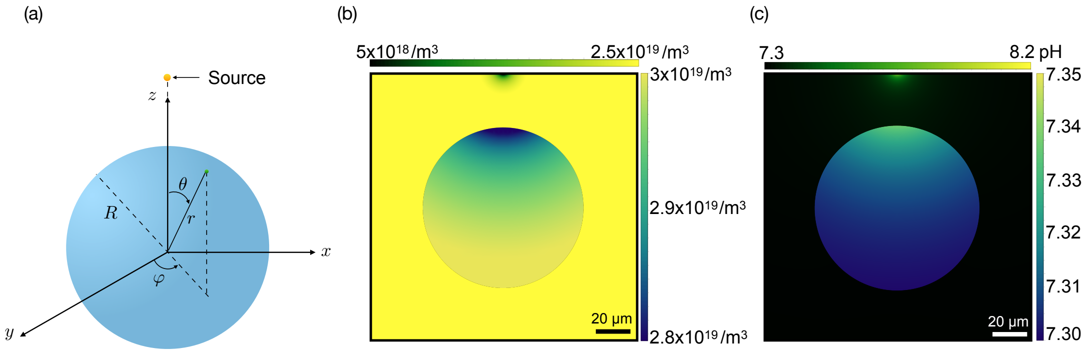


FIG. S18. (a) Schematic depiction of the system. (b) The concentration field inside and outside the droplet. (c) The variation of the pH value inside and outside the droplet. Note that in (b) and (c), different color schemes are used for inside and outside fields, with the color bar for the outside field shown on top.

equations as

$$\nabla^2 c = 0. \tag{22}$$

Note that here the advection of protons are negligible, as the relevant Pèclet number evaluates to $\mathrm{Pe} = \frac{VR}{D} \approx 10^{-4}$. The solution will be called $c^{\mathrm{in}}(r,\theta)$ for $r < R$ and $c^{\mathrm{out}}(r,\theta)$ for $r > R$, and they will be subject to the continuity of the concentration at the surface of the droplet as $c^{\mathrm{in}}\big|_{r=R} = c^{\mathrm{out}}\big|_{r=R}$. The continuity of the flux also demands $-D^{\mathrm{in}} \frac{\partial c^{\mathrm{in}}}{\partial r}\Big|_{r=R} = -D^{\mathrm{out}} \frac{\partial c^{\mathrm{out}}}{\partial r}\Big|_{r=R}$, where $D^{\mathrm{in}}$ and $D^{\mathrm{out}}$ are the diffusion coefficients inside and outside of the droplet, respectively. The solution for the concentration field inside and outside of the droplet is found in terms of a spherical coordinate set at the centre of the droplet as

$$c^{\mathrm{in}}(r,\theta) = c_\infty - \Delta c \left[\frac{R}{h} + \sum_{\ell=1} \frac{2\ell+1}{(1+\delta)\ell+1}\left(\frac{r}{h}\right)^{\ell} P_\ell(\cos\theta)\right] \tag{23}$$

$$c^{\mathrm{out}}(r,\theta) = c_\infty - \Delta c \sum_{\ell=1}\left[\frac{(1-\delta)\ell}{(1+\delta)\ell+1}\left(\frac{R^{2\ell+1}}{r^{\ell+1}h^{\ell}}\right) + \left(\frac{r}{h}\right)^{\ell}\right] P_\ell(\cos\theta) \tag{24}$$

where $\delta = D^{\mathrm{in}}/D^{\mathrm{out}}$ is the ratio between the diffusivities inside and outside, $\Delta c \equiv -I/(4\pi D^{\mathrm{out}} h)$ sets the scale of the concentration profiles in terms of the flux at the source (or sink), and $c_\infty$ is a constant that gives us the bulk concentration away from the droplet. Having $\Delta c$ at hand (see  for details on the numerical estimate), the concentration field can be illustrated for the inside and outside of the droplet, as shown in Fig. S18(b). We can alternatively show the variation of the pH value in the fluid; see Fig. S18(c).

*Flow field inside and outside of the droplet*

There are three mechanistically different contributions to the flow generated by the motion of the BSA proteins inside the droplet, and resultant flow in the medium: the *bulk diffusiophoretic flow* which stems from the high density of the proteins inside the droplet, the *Marangoni flow* due to the gradient of the surface tension, and the *surface diffusiophoretic flow* which gives rise to a tangential slip at the droplet-fluid interface. In what follows, we discuss each of these phenomena separately, and then provide the complete solution for the (inside and outside) flow field accounting for all three effects.

*Bulk Diffusiophoresis Effect* Due to the presence of the (proton) concentration gradient, the BSA proteins in the bulk of the droplet will experience a phoretic drift. While this interfacial transport phenomenon will normally lead to force-free drift in dilute concentrations away from boundaries, we expect the presence of the confining boundary of the droplet caused by the depletion forces of the polymers, the crowding effect of the proteins themselves inside the droplet, and the presence of the substrate to inflict opposing body forces on the BSA proteins across the droplet. These body forces will have a spatial distribution that will depend on these different considerations and contributions. Rather than resolving all these effects, we employ a simple description in which we assume these body forces are averaged out everywhere in the bulk of the droplet and look for the resulting flow profile that ensues from such a bulk-driven active droplet. Assuming a fluid-like behavior for the BSA proteins, we can write the Stokes equations for their motion as

$$-\eta^{\rm in}\nabla^2\boldsymbol{v}^{\rm in} = -\boldsymbol{\nabla}p^{\rm in} + \xi k_B T\boldsymbol{\nabla}c^{\rm in}, \tag{25}$$

where $\eta^{\rm in}$ is the viscosity inside the droplet, $k_B$ is the Boltzmann constant, $T$ is the temperature, and $\xi$ is a prefactor indicating the strength of the (spatially distributed) body force that results from the concentration gradient that drives the motion of the BSAs. For the flow outside the droplet, no additional body force contributes to the motion and we simply have

$$-\eta^{\rm out}\nabla^2\boldsymbol{v}^{\rm out} = -\boldsymbol{\nabla}p^{\rm out}. \tag{26}$$

Here we assume the BSAs are contained within the droplet, indicating that the radial component of the velocity is zero at the interface. We allow the tangential velocity to have a discontinuity at the interface, which due to surface friction, results in a discontinuity of the tangential shear stress as

$$\sigma_{r\theta}^{\rm in} - \sigma_{r\theta}^{\rm out} = \Gamma\left(v_\theta^{\rm in} - v_\theta^{\rm out}\right), \tag{27}$$

where $\Gamma$ is the friction coefficient for the surface slip.

*Marangoni Effect* Any gradient in the surface tension of the droplet can also lead to motion, both inside and outside the droplet. We have

$$\sigma_{r\theta}^{\rm in} - \sigma_{r\theta}^{\rm out} = -\left(\frac{\partial\gamma}{\partial c}\right)\frac{1}{R}\frac{\partial c^{\rm in}}{\partial\theta}, \tag{28}$$

where $\partial\gamma/\partial c$ indicates the rate at which the surface tension varies with changes in the concentration field. Combining this stress jump with the Stokes equations inside and outside the droplet, one can resolve the complete flow field induced by the Marangoni effect.

*Surface Diffusiophoresis Effect* In this case, instead of the shear stress, the tangential velocity has a discontinuity at the boundary, due to the diffusiophoretic effect. In particular, we have

$$v_\theta^{\rm in} - v_\theta^{\rm out} = \frac{\mu_s}{R}\frac{\partial c}{\partial\theta}, \tag{29}$$

where $\mu_s$ is the surface phoretic mobility.

*The Flow Profile* The Stokes equations need to be solved in combination with the incompressibility condition ($\boldsymbol{\nabla}\cdot\boldsymbol{v}^{\rm in} = 0$ and $\boldsymbol{\nabla}\cdot\boldsymbol{v}^{\rm out} = 0$), and the boundary condition that the normal (radial) component of the velocity vanishes at the droplet boundary. We do so for each of the above effects and (by relying on the linearity of the field equations) add their contributions together for the complete solution. Assuming the droplet is stationary, and that the BSAs are contained by the droplet interface, we can rely on the axisymmetric nature of the interactions and solve for the flow field using the Legendre expansion.

We define the viscosity ratio as $\Lambda = \eta^{\rm in}/\eta^{\rm out}$, and observe that the **Bulk** phoretic effect, the **Marangoni** effect, and the **Surface** phoretic effect, can be represented using the following velocity scales

$$V_{\rm B} = \frac{\xi k_B T R\Delta c}{\eta^{\rm in}}, \quad V_{\rm M} = \frac{\partial\gamma}{\partial c}\frac{\Delta c}{\eta^{\rm in}}, \quad V_{\rm S} = \frac{\Lambda\mu_s\Delta c}{R}, \tag{30}$$

respectively. Importantly, we observe that the three processes that lead to flow are characterized by different size dependencies: $V_{\rm B} \sim \Delta c R$, $V_{\rm M} \sim \Delta c R^0$, and $V_{\rm S} \sim \Delta c R^{-1}$. (Note that $\Delta c$ itself might have $R$ dependence but it would be the same for all three terms.) This implies that at the largest scales the bulk effect dominates, at the smallest scales the surface effect dominates and the Marangoni effect dominates at the intermediate sales.

The inner solution for the flow is found as

$$v_r^{\rm in}(r,\theta) = A_{\rm B}V_{\rm B} + A_{\rm M}V_{\rm M} + A_{\rm S}V_{\rm S}, \tag{31}$$

$$v_\theta^{\rm in}(r,\theta) = B_{\rm B}V_{\rm B} + B_{\rm M}V_{\rm M} + B_{\rm S}V_{\rm S}, \tag{32}$$

where the prefactors are geometrical functions given below

$$A_{\rm B} = \sum_{\ell=2}^{\infty} \frac{(\ell-1)(2\ell-1)}{2(2\ell+1)[\ell+\Lambda(\ell-1)]} \left(\frac{r^\ell}{h^{\ell-1}R}\right) \left[\left(\frac{R}{r}\right)^2 - 1\right] P_{\ell-1}(\cos\theta), \tag{33}$$

$$A_{\rm M} = \sum_{\ell=2}^{\infty} \frac{\ell(\ell-1)}{2(\Lambda+1)[\Lambda+(1+\Lambda)\ell]} \left(\frac{r^\ell}{h^{\ell-1}R}\right) \left[\left(\frac{R}{r}\right)^2 - 1\right] P_{\ell-1}(\cos\theta), \tag{34}$$

$$A_{\rm S} = -\sum_{\ell=2}^{\infty} \frac{\ell(\ell-1)(2\ell-1)}{(\Lambda+1)[\ell+\Lambda(\ell-1)]} \left(\frac{r^\ell}{h^{\ell-1}R}\right) \left[\left(\frac{R}{r}\right)^2 - 1\right] P_{\ell-1}(\cos\theta), \tag{35}$$

$$B_{\rm B} = \sum_{\ell=2}^{\infty} \frac{(\ell-1)}{2(2\ell+1)[\ell+\Lambda(\ell-1)]} \left(\frac{r^\ell}{h^{\ell-1}R}\right) \left[(\ell+2) - \ell\left(\frac{R}{r}\right)^2\right] \frac{P_{\ell-2}(\cos\theta) - P_\ell(\cos\theta)}{\sin\theta}, \tag{36}$$

$$B_{\rm M} = \sum_{\ell=2}^{\infty} \frac{\ell(\ell-1)}{2(\Lambda+1)(2\ell-1)[\Lambda+(1+\Lambda)\ell]} \left(\frac{r^\ell}{h^{\ell-1}R}\right) \left[(\ell+2) - \ell\left(\frac{R}{r}\right)^2\right] \frac{P_{\ell-2}(\cos\theta) - P_\ell(\cos\theta)}{\sin\theta}, \tag{37}$$

$$B_{\rm S} = -\sum_{\ell=2}^{\infty} \frac{\ell(\ell-1)}{(\Lambda+1)[\ell+\Lambda(\ell-1)]} \left(\frac{r^\ell}{h^{\ell-1}R}\right) \left[(\ell+2) - \ell\left(\frac{R}{r}\right)^2\right] \frac{P_{\ell-2}(\cos\theta) - P_\ell(\cos\theta)}{\sin\theta}. \tag{38}$$

Note that $v_r^{\rm in}(r=R,\theta)$ vanishes identically but $v_\theta^{\rm in}(r=R,\theta)$ remains finite.

For the flow field outside of the droplet we find

$$v_r^{\rm out}(r,\theta) = M_{\rm B}V_{\rm B} + M_{\rm M}V_{\rm M} + \frac{1}{\Lambda}M_{\rm S}V_{\rm S}, \tag{39}$$

$$v_\theta^{\rm out}(r,\theta) = N_{\rm B}V_{\rm B} + N_{\rm M}V_{\rm M} + \frac{1}{\Lambda}N_{\rm S}V_{\rm S}, \tag{40}$$

where

$$M_{\rm B} = \sum_{\ell=2}^{\infty} \frac{(\ell-1)(2\ell-1)}{2(2\ell+1)[\ell+(\ell-1)\Lambda]} \left[\frac{(2\ell-1) - R\Gamma/\eta_{\rm in}}{(2\ell-1)\Lambda + R\Gamma/\eta_{\rm in}}\right] \left(\frac{R^{2\ell-2}}{r^{\ell-1}h^{\ell-1}}\right) \left[\left(\frac{R}{r}\right)^2 - 1\right] P_{\ell-1}(\cos\theta) \tag{41}$$

$$M_{\rm M} = -\sum_{\ell=2}^{\infty} \frac{\ell(\ell-1)}{2(1+\Lambda)[\Lambda+(1+\Lambda)\ell]} \left(\frac{R^{2\ell-2}}{r^{\ell-1}h^{\ell-1}}\right) \left[\left(\frac{R}{r}\right)^2 - 1\right] P_{\ell-1}(\cos\theta), \tag{42}$$

$$M_{\rm S} = \sum_{\ell=2}^{\infty} \frac{\ell(\ell-1)(2\ell-1)}{2(\Lambda+1)[\ell+\Lambda(\ell-1)]} \left(\frac{R^{2\ell-2}}{r^{\ell-1}h^{\ell-1}}\right) \left[\left(\frac{R}{r}\right)^2 - 1\right] P_{\ell-1}(\cos\theta), \tag{43}$$

$$N_{\rm B} = -\sum_{\ell=2}^{\infty} \frac{(\ell-1)}{2(2\ell+1)[\ell+(\ell-1)\Lambda]} \left[\frac{(2\ell-1) - R\Gamma/\eta_{\rm in}}{(2\ell-1)\Lambda + R\Gamma/\eta_{\rm in}}\right] \left[(\ell-3) - (\ell-1)\left(\frac{R}{r}\right)^2\right] \frac{P_{\ell-2}(\cos\theta) - P_\ell(\cos\theta)}{\sin\theta}, \tag{44}$$

$$N_{\rm M} = \sum_{\ell=2}^{\infty} \frac{\ell(\ell-1)}{2(1+\Lambda)(2\ell-1)[\Lambda+(1+\Lambda)\ell]} \left(\frac{R^{2\ell-2}}{r^{\ell-1}h^{\ell-1}}\right) \left[(\ell-3) - (\ell-1)\left(\frac{R}{r}\right)^2\right] \frac{P_{\ell-2}(\cos\theta) - P_\ell(\cos\theta)}{\sin\theta}, \tag{45}$$

$$N_{\rm S} = -\sum_{\ell=2}^{\infty} \frac{\ell(\ell-1)}{2(\Lambda+1)[\ell+\Lambda(\ell-1)]} \left(\frac{R^{2\ell-2}}{r^{\ell-1}h^{\ell-1}}\right) \left[(\ell-3) - (\ell-1)\left(\frac{R}{r}\right)^2\right] \frac{P_{\ell-2}(\cos\theta) - P_\ell(\cos\theta)}{\sin\theta}, \tag{46}$$

are the corresponding geometrical functions.

#### *Estimating the Parameters*

Using the values from the experiments we have $R = 90\,\mu$m, $h = 155\,\mu$m, and $\delta \approx 1.4 \times 10^{-2}$. While we do not know the flux at the source $I$, we can calculate it by using the pH values at two points, namely the pipette and far from it in the buffer, as well as the size of the pipette $r_{\rm p} = 5\,\mu$m. We find

$$\Delta c = \frac{\left(10^{-{\rm pH}_{\rm buffer}+3} - 10^{-{\rm pH}_{\rm pipette}+3}\right) N_{\rm Av}}{\sum_{\ell=1}\left[\frac{(1-\delta)\ell}{(1+\delta)\ell+1}\left(\frac{R}{h}\right)^{2l+1} + \left(1-\frac{r_{\rm p}}{h}\right)^{\ell}\right]} \approx 8.7 \times 10^{17}/{\rm m}^3, \tag{47}$$

where $N_{\mathrm{Av}} = 6.02 \times 10^{23}$ is the Avogadro number, $\mathrm{pH_{pipette}} = 8.2$ and $\mathrm{pH_{buffer}} = 7.3$.

We can make an estimate for the parameter as $\xi \approx \frac{\eta^{\mathrm{in}}}{\eta^{\mathrm{out}}} \frac{\lambda^2}{a_{\mathrm{BSA}}^2}$, where $\lambda$ is the Debye length, and $a_{\mathrm{BSA}}$ is the BSA radius. We define the Bjerrum length as $\ell_B = e^2/(\varepsilon k_B T)$, where $e$ is the electronic charge and $\varepsilon$ is the permittivity of free space. The Debye length $\lambda$ is then defined via $\lambda^{-2} = 8\pi\ell_B \sum_i z_i^2 c_i$, where $z_i$ is the valence of species $i$ with concentration $c_i$. Using $a_{\mathrm{BSA}} = 4$ nm, and $\lambda = 5$ nm, which results from reasonable values for the ionic strength of the buffer, we find that our rough estimate gives $\xi \approx 100$, which is consistent with the observations. Using this estimate, we obtain $V_{\mathrm{B}} \approx 16$ $\mu$m/s.

For the surface-slip friction coefficient, we can use an estimate of $\Gamma \approx \eta^{\mathrm{out}}/L_{\mathrm{slip}}$, where the slip length of the dense protein droplet $L_{\mathrm{slip}}$ can be estimated as a protein radius, namely $L_{\mathrm{slip}} \approx a_{\mathrm{BSA}}$. Therefore, $\Gamma \approx \eta^{\mathrm{out}}/a_{\mathrm{BSA}}$. Using $\eta^{\mathrm{out}} = 0.012$ Pa.s and $a_{\mathrm{BSA}} = 4$ nm, we obtain $\Gamma \approx 10^7$ N.s/m$^3$.

For the Marangoni velocity scale, using $\partial\gamma/\partial c = 2 \times 10^{-23}$ N/m$^4$ we obtain $V_{\mathrm{M}} \approx 8$ $\mu$m/s. Therefore, it appears that for the sizes of droplets in our experiments the bulk flow and the Marangoni flow are in direction competition.

The surface phoretic mobility can be estimated as $\mu_s \approx k_B T \lambda^2/\eta_{\mathrm{in}}$. By taking $k_B T = 4.11 \times 10^{-21} J$, $\eta_{\mathrm{in}} = 2$ Pa.s, $\lambda = 5$ nm, we find $\mu_s \approx 10^{-37}$ m$^5$/s. Using this estimate, we obtain $V_{\mathrm{S}} \approx 10^{-11}$ $\mu$m/s, which shows that for our experiments the surface phoretic flow can be safely neglected.

---

* These authors contributed equally.
†
‡

[1] S. B. Zimmerman and S. O. Trach, Journal of molecular biology **222**, 599 (1991).
[2] J. Van Den Berg, A. J. Boersma, and B. Poolman, Nature Reviews Microbiology **15**, 309 (2017).
[3] A. J. Boersma, I. S. Zuhorn, and B. Poolman, Nature methods **12**, 227 (2015).
[4] M. C. Konopka, K. A. Sochacki, B. P. Bratton, I. A. Shkel, M. T. Record, and J. C. Weisshaar, Journal of bacteriology **191**, 231 (2009).
[5] S. Cayley and M. T. Record Jr, Journal of Molecular Recognition **17**, 488 (2004).
[6] G.-W. Li, D. Burkhardt, C. Gross, and J. S. Weissman, Cell **157**, 624 (2014).
[7] K. Valgepea, K. Adamberg, A. Seiman, and R. Vilu, Molecular BioSystems **9**, 2344 (2013).
[8] P. Lu, C. Vogel, R. Wang, X. Yao, and E. M. Marcotte, Nature biotechnology **25**, 117 (2007).
[9] L. Arike, K. Valgepea, L. Peil, R. Nahku, K. Adamberg, and R. Vilu, Journal of proteomics **75**, 5437 (2012).
[10] H.-X. Zhou, G. Rivas, and A. P. Minton, Annu. Rev. Biophys. **37**, 375 (2008).
[11] W. M. Aumiller Jr, B. W. Davis, E. Hatzakis, and C. D. Keating, The Journal of Physical Chemistry B **118**, 10624 (2014).
[12] S. F. Banani, H. O. Lee, A. A. Hyman, and M. K. Rosen, Nature reviews Molecular cell biology **18**, 285 (2017).
[13] M. Boehning, C. Dugast-Darzacq, M. Rankovic, A. S. Hansen, T. Yu, H. Marie-Nelly, D. T. McSwiggen, G. Kokic, G. M. Dailey, P. Cramer, *et al.*, Nature structural & molecular biology **25**, 833 (2018).
[14] W. J. Altenburg, N. A. Yewdall, D. F. Vervoort, M. H. Van Stevendaal, A. F. Mason, and J. C. van Hest, Nature communications **11**, 1 (2020).
[15] S. Boeynaems, S. Alberti, N. L. Fawzi, T. Mittag, M. Polymenidou, F. Rousseau, J. Schymkowitz, J. Shorter, B. Wolozin, L. Van Den Bosch, *et al.*, Trends in cell biology **28**, 420 (2018).
[16] A. A. Hyman, C. A. Weber, and F. Jülicher, Annual review of cell and developmental biology **30**, 39 (2014).
[17] C. P. Brangwynne, C. R. Eckmann, D. S. Courson, A. Rybarska, C. Hoege, J. Gharakhani, F. Jülicher, and A. A. Hyman, Science **324**, 1729 (2009).
[18] M. Feric, N. Vaidya, T. S. Harmon, D. M. Mitrea, L. Zhu, T. M. Richardson, R. W. Kriwacki, R. V. Pappu, and C. P. Brangwynne, Cell **165**, 1686 (2016).
[19] S. Boeynaems, A. S. Holehouse, V. Weinhardt, D. Kovacs, J. Van Lindt, C. Larabell, L. Van Den Bosch, R. Das, P. S. Tompa, R. V. Pappu, *et al.*, Proceedings of the National Academy of Sciences **116**, 7889 (2019).
[20] P. M. McCall, S. Srivastava, S. L. Perry, D. R. Kovar, M. L. Gardel, and M. V. Tirrell, Biophysical journal **114**, 1636 (2018).
[21] L. Faltova, A. M. Küffner, M. Hondele, K. Weis, and P. Arosio, ACS nano **12**, 9991 (2018).
[22] R. R. Poudyal, R. M. Guth-Metzler, A. J. Veenis, E. A. Frankel, C. D. Keating, and P. C. Bevilacqua, Nature communications **10**, 1 (2019).
[23] H. Karoui, M. J. Seck, and N. Martin, Chemical Science **12**, 2794 (2021).
[24] W. M. Aumiller and C. D. Keating, Nature chemistry **8**, 129 (2016).
[25] K. K. Nakashima, J. F. Baaij, and E. Spruijt, Soft Matter **14**, 361 (2018).
[26] S. Deshpande, F. Brandenburg, A. Lau, M. G. Last, W. K. Spoelstra, L. Reese, S. Wunnava, M. Dogterom, and C. Dekker, Nature communications **10**, 1 (2019).
[27] C. Donau, F. Späth, M. Sosson, B. A. Kriebisch, F. Schnitter, M. Tena-Solsona, H.-S. Kang, E. Salibi, M. Sattler, H. Mutschler, *et al.*, Nature communications **11**, 1 (2020).
[28] Z. W. Lim, Y. Ping, and A. Miserez, Bioconjugate chemistry **29**, 2176 (2018).
[29] R. A. Copeland, *Enzymes: a practical introduction to structure, mechanism, and data analysis* (John Wiley & Sons, 2000).
[30] K. A. Johnson and R. S. Goody, Biochemistry **50**, 8264 (2011).
[31] G. Johansson, Journal of Chromatography B: Biomedical Sciences and Applications **680**, 123 (1996), 9th International Conference On Partitioning in Aqueous Two-

Phase Systems.
[32] E. R. Dufresne, arXiv preprint arXiv:1903.09584 (2019).
[33] R. Golestanian, Phys. Rev. Lett. **115**, 108102 (2015).
[34] R. Milo, BioEssays **35**, 1050 (2013).
[35] R. J. Ellis, Trends in biochemical sciences **26**, 597 (2001).
[36] R. Milo and R. Phillips, *Cell biology by the numbers* (Garland Science, 2015).
[37] D. S. Goodsell, Trends in biochemical sciences **16**, 203 (1991).
[38] K. Nishizawa, K. Fujiwara, M. Ikenaga, N. Nakajo, M. Yanagisawa, and D. Mizuno, Scientific reports **7**, 1 (2017).
[39] M. Guo, A. J. Ehrlicher, M. H. Jensen, M. Renz, J. R. Moore, R. D. Goldman, J. Lippincott-Schwartz, F. C. Mackintosh, and D. A. Weitz, Cell **158**, 822 (2014).
[40] R. J. Gay, R. B. McComb, and G. N. Bowers Jr, Clinical chemistry **14**, 740 (1968).
[41] E. Jackson, F. López-Gallego, J. Guisan, and L. Betancor, Process Biochemistry **51**, 1248 (2016).
[42] M. W. Eggert, M. E. Byrne, and R. P. Chambers, Applied biochemistry and biotechnology **165**, 676 (2011).
[43] R. Stambaugh and D. Post, Journal of Biological Chemistry **241**, 1462 (1966).
[44] A. M. Makarieva, V. G. Gorshkov, B.-L. Li, S. L. Chown, P. B. Reich, and V. M. Gavrilov, Proceedings of the National Academy of Sciences **105**, 16994 (2008).
[45] K. Kappaun, A. R. Piovesan, C. R. Carlini, and R. Ligabue-Braun, Journal of advanced research **13**, 3 (2018).
[46] C. Riedel, R. Gabizon, C. A. Wilson, K. Hamadani, K. Tsekouras, S. Marqusee, S. Pressé, and C. Bustamante, Nature **517**, 227 (2015).
[47] M. Ijavi, R. W. Style, L. Emmanouilidis, A. Kumar, S. M. Meier, A. L. Torzynski, F. H. Allain, Y. Barral, M. O. Steinmetz, and E. R. Dufresne, Soft Matter **17**, 1655 (2021).
[48] D. B. Stein, G. De Canio, E. Lauga, M. J. Shelley, and R. E. Goldstein, Physical Review Letters **126**, 028103 (2021).
[49] L. Scriven and C. Sternling, Nature **187**, 186 (1960).
[50] D. J. Bonthuis and R. Golestanian, Phys. Rev. Lett. **113**, 148101 (2014).
[51] B. Ramm, A. Goychuk, A. Khmelinskaia, P. Blumhardt, H. Eto, K. A. Ganzinger, E. Frey, and P. Schwille, Nature Physics (2021), 10.1038/s41567-021-01213-3.
[52] R. Golestanian, arXiv:1909.03747 (2019), arXiv:1909.03747.
[53] N. Lane, BioEssays **39**, 1600217 (2017).
[54] C. Bonfio, E. Godino, M. Corsini, F. F. de Biani, G. Guella, and S. S. Mansy, Nature Catalysis **1**, 616 (2018).
[55] W. F. Martin, F. L. Sousa, and N. Lane, science **344**, 1092 (2014).
[56] R. E. Goldstein and J.-W. van de Meent, Interface focus **5**, 20150030 (2015).
[57] T. Shimmen and E. Yokota, Current opinion in cell biology **16**, 68 (2004).
[58] K. Luby-Phelps, *Microcompartmentation and Phase Separation in Cytoplasm*, International Review of Cytology, **192**, 189 (1999).
[59] H.-X. Zhou, Archives of Biochemistry and Biophysics **469**, 76 (2008).
[60] A. P. Minton, Journal of biological chemistry **276**, 10577 (2001).
[61] A. P. Minton, Journal of pharmaceutical sciences **94**, 1668 (2005).
[62] A. Zecchin, P. C. Stapor, J. Goveia, and P. Carmeliet, Current opinion in biotechnology **34**, 73 (2015).
[63] A. H. Chen and P. A. Silver, Trends in cell biology **22**, 662 (2012).
[64] U. Heinig, M. Gutensohn, N. Dudareva, and A. Aharoni, Current opinion in biotechnology **24**, 239 (2013).
[65] B. P. Tu, A. Kudlicki, M. Rowicka, and S. L. McKnight, Science **310**, 1152 (2005).
[66] C. Schmid-Dannert and F. López-Gallego, Current opinion in chemical biology **49**, 97 (2019).
[67] S. Sengupta, K. K. Dey, H. S. Muddana, T. Tabouillot, M. E. Ibele, P. J. Butler, and A. Sen, Journal of the American Chemical Society **135**, 1406 (2013).
[68] A.-Y. Jee, S. Dutta, Y.-K. Cho, T. Tlusty, and S. Granick, Proc. Natl. Acad. Sci. U. S. A. **115**, 14 (2018).
[69] J. Agudo-Canalejo, P. Illien, and R. Golestanian, Nano Lett. **18**, 2711 (2018).
[70] R. Golestanian, Nature Physics **13**, 323 (2016).
[71] D. Zwicker, R. Seyboldt, C. A. Weber, A. A. Hyman, and F. Jülicher, Nature Physics **13**, 408 (2017).
[72] A. I. Oparin *et al.*, The origin of life on the earth. (1957).
[73] T. Z. Jia, K. Chandru, Y. Hongo, R. Afrin, T. Usui, K. Myojo, and H. J. Cleaves, Proceedings of the National Academy of Sciences **116**, 15830 (2019).
[74] J. W. Szostak, D. P. Bartel, and P. L. Luisi, Nature **409**, 387 (2001).
[75] A. J. Dzieciol and S. Mann, Chemical Society Reviews **41**, 79 (2012).
[76] T. D. Tang, C. R. C. Hak, A. J. Thompson, M. K. Kuimova, D. Williams, A. W. Perriman, and S. Mann, Nature chemistry **6**, 527 (2014).
[77] R. R. Poudyal, F. Pir Cakmak, C. D. Keating, and P. C. Bevilacqua, Biochemistry **57**, 2509 (2018).
[78] Y. Qiao, M. Li, R. Booth, and S. Mann, Nature chemistry **9**, 110 (2017).
[79] B. P. Kumar, A. J. Patil, and S. Mann, Nature chemistry **10**, 1154 (2018).
[80] A. Joesaar, S. Yang, B. Bögels, A. van der Linden, P. Pieters, B. P. Kumar, N. Dalchau, A. Phillips, S. Mann, and T. F. de Greef, Nature nanotechnology **14**, 369 (2019).
[81] P. Gobbo, A. J. Patil, M. Li, R. Harniman, W. H. Briscoe, and S. Mann, Nature materials **17**, 1145 (2018).
[82] R. Booth, Y. Qiao, M. Li, and S. Mann, Angewandte Chemie **131**, 9218 (2019).
[83] T. Lu and E. Spruijt, Journal of the American Chemical Society **142**, 2905 (2020).
[84] E. Sokolova, E. Spruijt, M. M. Hansen, E. Dubuc, J. Groen, V. Chokkalingam, A. Piruska, H. A. Heus, and W. T. Huck, Proceedings of the National Academy of Sciences **110**, 11692 (2013).
[85] F. Spaeth, C. Donau, A. M. Bergmann, M. Kränzlein, C. V. Synatschke, B. Rieger, and J. Boekhoven, Journal of the American Chemical Society **143**, 4782 (2021).
[86] R. D. Alvares, A. Hasabnis, R. S. Prosser, and P. M. Macdonald, Analytical chemistry **88**, 3730 (2016).
[87] H. T. Spanke, R. W. Style, C. François-Martin, M. Feofilova, M. Eisentraut, H. Kress, J. Agudo-Canalejo, and E. R. Dufresne, Physical Review Letters **125**, 198102 (2020).
[88] S. Minikami and C. de Verdier, Eur J Biochem **65**, 451 (1976).
